# OMEGA: a software tool for the management, analysis, and dissemination of intracellular trafficking data that incorporates motion type classification and quality control

**DOI:** 10.1101/251850

**Authors:** Alessandro Rigano, Vanni Galli, Jasmine M. Clark, Lara E. Pereira, Loris Grossi, Jeremy Luban, Raffaello Giulietti, Tiziano Leidi, Eric Hunter, Mario Valle, Ivo F. Sbalzarini, Caterina Strambio-De-Castillia

**Affiliations:** Program In Molecular Medicine, University of Massachusetts Medical School, Worcester, Massachusetts 01605, United States of America; Istituto Sistemi Informativi e Networking, Scuola Universitaria Professionale della Svizzera Italiana, CH-6928 Manno, Switzerland; Yerkes National Primate Research Center, Emory University, Atlanta, Georgia, 30329, United States of America; CSCS - Swiss National Supercomputing Centre, ETH-Zurich, CH-6900 Lugano, Switzerland; MOSAIC Group, Center for Systems Biology Dresden; TU Dresden, Faculty of Computer Science; Max Planck Institute of Molecular Cell Biology and Genetics, Pfotenhauerstr. 108, 01307 Dresden, Germany; Emory University Nell Hodgson Woodruff School of Nursing, Atlanta, Georgia, 30322, United States of America

## Abstract

**MOTIVATION:** Particle tracking coupled with time-lapse microscopy is critical for understanding the dynamics of intracellular processes of clinical importance. Spurred on by advances in the spatiotemporal resolution of microscopy and automated computational methods, this field is increasingly amenable to multi-dimensional high-throughput data collection schemes (Snijder et al., 2012). Typically, complex particle tracking datasets generated by individual laboratories are produced with incompatible methodologies that preclude comparison to each other. There is therefore an unmet need for data management systems that facilitate data standardization, meta-analysis, and structured data dissemination. The integration of analysis, visualization, and quality control capabilities into such systems would eliminate the need for manual transfer of data to diverse downstream analysis tools. At the same time, it would lay the foundation for shared trajectory data, particle tracking, and motion analysis standards.

**RESULTS:** Here, we present Open Microscopy Environment inteGrated Analysis (OMEGA), a cross-platform data management, analysis, and visualization system, for particle tracking data, with particular emphasis on results from viral and vesicular trafficking experiments. OMEGA provides intuitive graphical interfaces to implement integrated particle tracking and motion analysis workflows while providing easy to use facilities to automatically keep track of error propagation, harvest data provenance and ensure the persistence of analysis results and metadata. Specifically, OMEGA: 1) imports image data and metadata from data management tools such as the Open Microscopy Environment Remote Objects (OMERO; Allan et al., 2012); 2) tracks intracellular particles movement; 3) facilitates parameter optimization and trajectory results inspection and validation; 4) performs downstream trajectory analysis and motion type classification; 5) estimates the uncertainty propagating through the motion analysis pipeline; and, 6) facilitates storage and dissemination of analysis results, and analysis definition metadata, on the basis of our newly proposed FAIRsharing.org complainant Minimum Information About Particle Tracking Experiments (MIAPTE; Rigano and Strambio-De-Castillia, 2016; 2017) guidelines in combination with the OME-XML data model (Goldberg et al., 2005). In so doing, OMEGA maintains a persistent link between raw image data, intermediate analysis steps, the overall analysis output, and all necessary metadata to repeat the analysis process and reproduce its results.

**Availability and implementation:** OMEGA is a cross-platform, open-source software developed in Java. Source code and cross-platform binaries are freely available on GitHub at https://github.com/OmegaProject/Omega (doi: 10.5281/zenodo.2535523), under the GNU General Public License v.3.

**Supplementary information:** Supplementary Material is available at BioRxiv.org

## 1 Introduction

### 1.1 Description of the problem

Dynamic intracellular processes, such as viral and bacterial infection (Chenouard et al., 2009a; Brandenburg and Zhuang, 2007; Sun et al., 2013; Mercer et al., 2010; Li et al., 2017; Pereira et al., 2014; Ewers et al., 2005), vesicular trafficking (Siebrasse et al., 2016; Aoyama et al., 2017; Gramlich and Klyachko, 2017; Jandt et al., 2011; Jandt and Zeng, 2012; Loerke et al., 2009), membrane receptors dynamics (Jaqaman et al., 2016; Sergé et al., 2008; Block et al., 2016; Saxton and Jacobson, 1997), cytoskeletal rearrangement (Applegate et al., 2011; Akhmanova and Steinmetz, 2008), focal adhesion reorganization (Berginski et al., 2011), gene transcription (Sinha et al., 2010), and genome maintenance (Agarwal et al., 2011) are important for many clinically relevant fields of study including immune regulation, metabolic disorders, infectious diseases, and cancer. In all of these cases, diverse individual sub-resolution ‘particles’ (e.g. single molecules, microtubule tips, viruses, vesicles and organelles) dynamically interact with a large number of cellular structures that influence trajectory and speed. The path followed by individual particles varies significantly depending on molecular composition, cargo, and destination. Given the changing and multi-step nature of several of these processes, many questions would benefit from studying them in living cells. The fundamental spatial and temporal heterogeneity of these trafficking processes emphasizes the importance of utilizing single-particle measurements, rather than ensemble averages or flow measurements, in order to gain insight into molecular mechanisms, predict outcome, and rationally design effective therapeutic interventions. The time-resolved visualization of individual heterogeneous intracellular particles by fluorescence-microscopy, coupled with feature point tracking techniques - referred to as Single-Particle Tracking (SPT) (De Brabander et al., 1985) or Multiple-Particle Tracking (MPT) (Genovesio et al., 2006) - and mathematical analysis of motion, is ideally suited to follow the fate of particles as they progress within the cell, to map fleeting interactions with other cellular components, and to dissect individual transport steps. For example, single viral imaging experiments coupled with SPT have improved understanding of the early phases of viral entry and revealed previously un-recognized entry stages (Brandenburg and Zhuang, 2007; Flatt and Greber, 2017; Sun et al., 2013; Greber and Way, 2006; Wang et al., 2017; Ewers et al., 2005; Helmuth et al., 2007; Yamauchi et al., 2011).

As a consequence of the steady improvement of the spatiotemporal resolution of microscopic techniques, advances in automated particle tracking and motion analysis (Sbalzarini and Koumoutsakos, 2005; Arhel et al., 2006; Jaqaman et al., 2008; Chenouard et al., 2014; Smith et al., 2015; Genovesio et al., 2006), and the availability of software tools (Carpenter et al., 2012; Schindelin et al., 2012; de Chaumont et al., 2012; Eliceiri et al., 2012; Perry et al., 2012; Swedlow and Eliceiri, 2009; Tinevez et al., 2016; Jaqaman et al., 2008; Kalaidzidis, 2009; Incardona and Sbalzarini, 2014), SPT holds the promise of becoming amenable to multi-dimensional high-throughput data collection schemas (Damm and Pelkmans, 2006; Snijder et al., 2012; Rämö et al., 2014; Taute et al., 2015).

However, many of the fundamental image data management limitations holding back “[…] the routine application of automated image analysis […] to large volumes of information generated by digital imaging” (verbatim from: Swedlow et al., 2003) are still in place even several years after the initial identification of the problem. As a case in point, the utilization of viral particle tracking to draw direct real-time correlations between alterations in viral mobility and underlying perturbations in the viral and cellular states, remains a considerable challenge even at low-throughput, and it is difficult if not impossible to scale to the systems biology level (Arhel et al., 2006; McDonald et al., 2002; Mamede et al., 2017; Mamede and Hope, 2016; Sood et al., 2017). As a result, most virology studies to date rely on biochemical and genetic analyses conducted in bulk and on the microscopic analysis of fixed cells, which fail to capture viral heterogeneity and the complexity of viral infection processes. Moreover, when particle tracking data is obtained, the datasets produced at individual laboratories are difficult to compare with analogous data generated at other times or places, making integration with different data types, and meta-analysis, impossible.

This situation reveals the presence of an unmet need for tools that allow the management of large datasets of intracellular trafficking data in a manner that would streamline analysis workflows, unify dataflow, automate the harvesting of data provenance, facilitate results inspection, quality control and uncertainty estimation, foster reproducibility, dissemination and meta-analysis, and, ultimately, lay the foundation for the creation of distributed particle tracking data commons.

### 1.2 Statement of purpose: integration of particle tracking work- and dataflows with automated data provenance harvesting and uncertainty estimation

A major hurdle preventing particle tracking from becoming a routine high-throughput cell biology technique, the results of which can be reproduced and compared across different data production sites, is related to the size and complexity of the data. The number of particles within each cell may be in the hundreds, images typically contain MBs of data, experiment may produce thousands of images and the correct interpretation of results depends on the knowledge of the experimental, optical and image-analysis context. Hence it follows that in order to tackle this problem, automated image acquisition, processing, and analysis have to be closely coupled with robust and standardized data management methods, and with accurate accounting of error propagation and data provenance. Although software tools exist to execute several steps of the particle tracking workflow, tools for the integrated and automated execution of key data provenance harvesting, data management and uncertainty evaluation steps do not exist (Table I) (Tinevez et al., 2016; de Chaumont et al., 2012). The development of user-oriented, freely available, integrated data and metadata management and analysis systems for particle tracking data would thus offer a timely next step for the field, as the benefit of this approach has been well established in other fields, where data management and dissemination infrastructure is more mature (Data models to GO-FAIR., 2017; Wilkinson et al., 2016; UniProt Consortium, 2015; Benson et al., 2012; Berman et al., 2003; Wenger et al., 2000).

**Table I.**
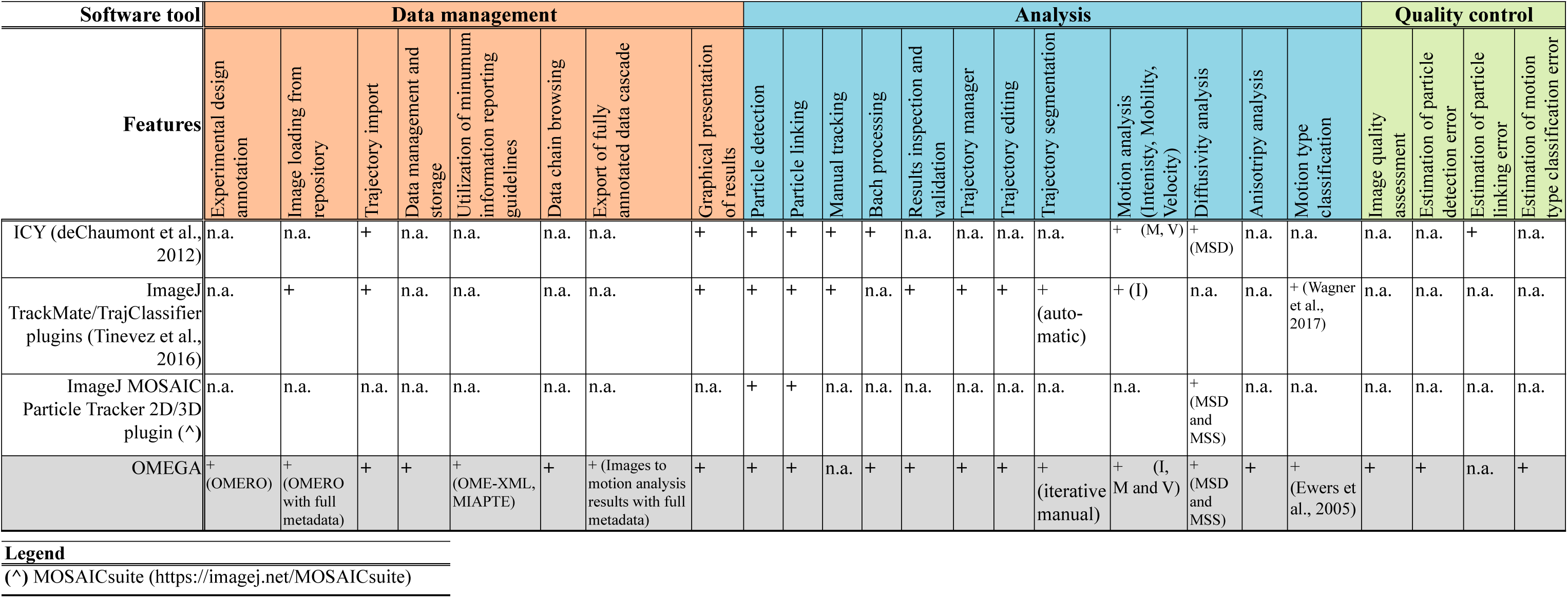
Overview of currently available integrated software tools supporting particle tracking and motion analysis workflows

In order to address this difficulty, we introduce a novel cross-platform, open-source software called Open Microscopy Environment inteGrated Analysis (OMEGA; Figure 1), which provides a rich graphical user interface (GUI; Figure 2; Supplemental Information 1) to aid the user with particle tracking data production, analysis, validation, uncertainty estimation visualization, and management.

**Figure 1:**
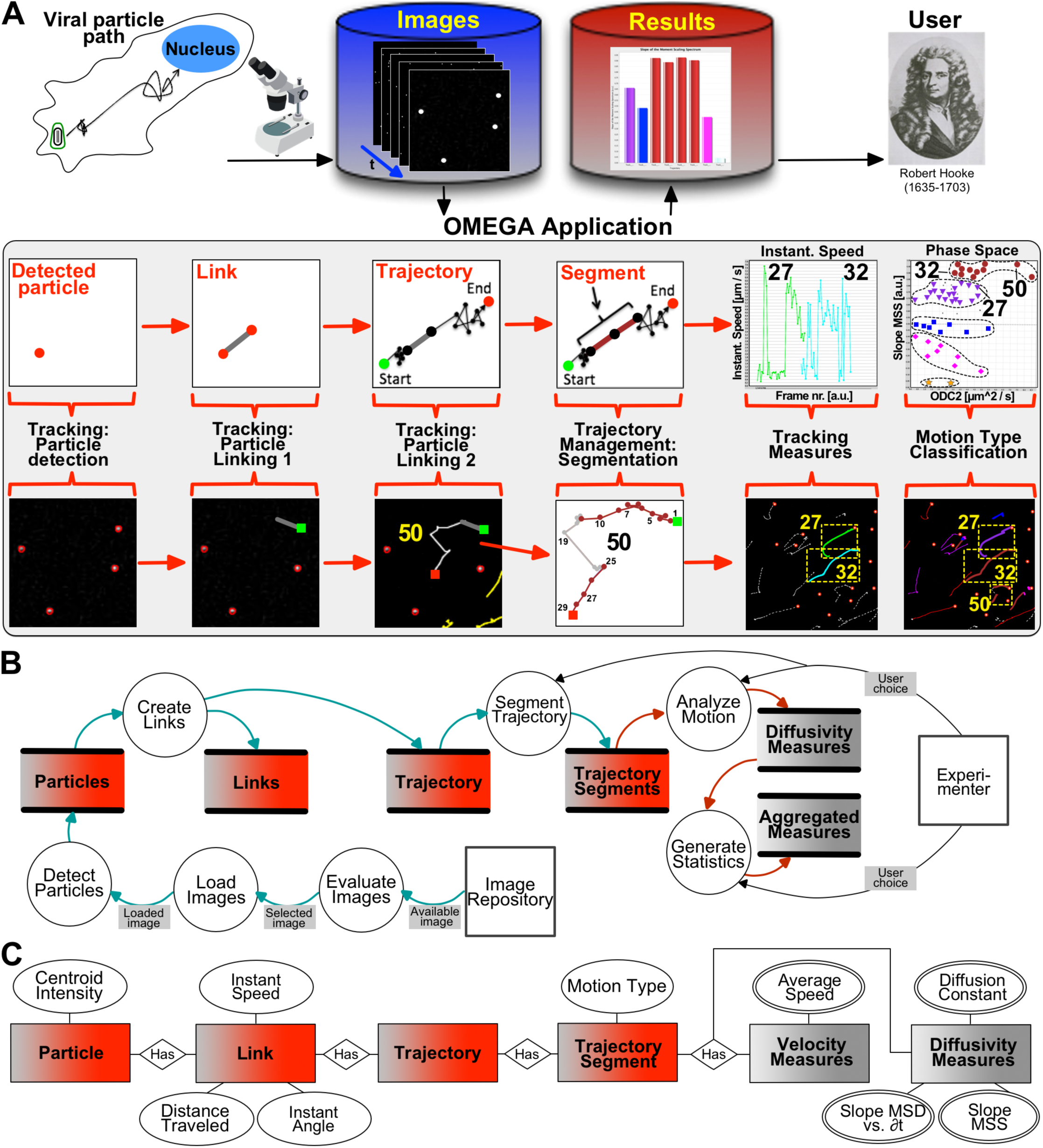
End-to-end, automatic reproducibility and repeatability support for particle tracking and motion analysis experiments in OMEGA on the basis of a MIAPTE compliant data model. OMEGA provides end-to-end supports for the reproducibility and repeatability of particle tracking and motion analysis experiments by providing an integrated platform that: 1) executes all basic steps of a particle tracking and motion analysis steps workflow. 2) Tracks uncertainty propagation across the analysis pipeline. 3) Automatically maintains persistent links between the chain of data, harvests data provenance and utilizes it to annotate analysis result. 4) Utilizes a domain-relevant community standards to describe data, metadata and processing steps. Depicted here is a schematic diagram representing the system context in which OMEGA operates, and the workflow (A) required for the estimation of the sub-cellular trajectories followed by diffraction-limited, intra-cellular viral particles, and the computation of biologically meaningful measures from particles coordinates. A) Images are acquired using available microscopes, and imported into available OMERO databases. Subsequently, images are imported into OMEGA using the Image Data Browser plugin, and subjected to Single Particle Tracking (SPT) in two independent steps using plugins implementing the Particle Tracking module (i.e. two separate Particle Detection and Particle Linking plugins, or a single unified Particle Tracking plugin). As needed individual trajectories can be subdivided into uniform segments using the interactive Trajectory Segmentation plugin. In the example, trajectory nr. 50 was subdivided into three segments, two of which were assigned the Directed motion type (*maroon*) and the third one was left unclassified (*grey*). In addition, all other trajectories, which appeared to be uniform in nature were assigned the color corresponding to the predicted motion type depending on the observed slope of the MSS curve (*grey*, not assigned; *yellow*, confined; fuchsia, sub-diffusive; *blue*, diffusive; *purple*, super-diffusive; *maroon*, directed). Trajectories were then subjected to motion analysis using the Velocity Tracking Measures (VTM) and the Diffusivity Tracking Measures (DTM) plugins. Instantaneous Speed results for trajectory nr. 27 and 32, and D vs. SMSS Phase space results for all trajectories are displayed. The position of spots representing trajectory nr. 27, 32 and 50 are indicated. B) The path taken by data across the workflow indicated in panel A is represented using the data flow diagram (DFD) formalism. Circles represent processes that transform data, arrows represent data in motion, arrow labels represent specific packages of data being moved, double lined-rectangles represent data at rest (i.e. data stores) and squares represent entities (i.e. Experimenter(s) and external data repositories) that interact with the system from the outside. C) In order to ensure the preservation of data provenance links, OMEGA utilizes data structures whose model in based on our recently proposed Minimum Information About Particle Tracking (MIAPTE) guidelines. Depicted here is an Entity Relationship diagram representing the corresponding OME-XML (*blue*) MIAPTE (*red*, particle data; *grey*, analysis data) elements utilized to capture data pertaining to each step of the data-flow.

**Figure 2:**
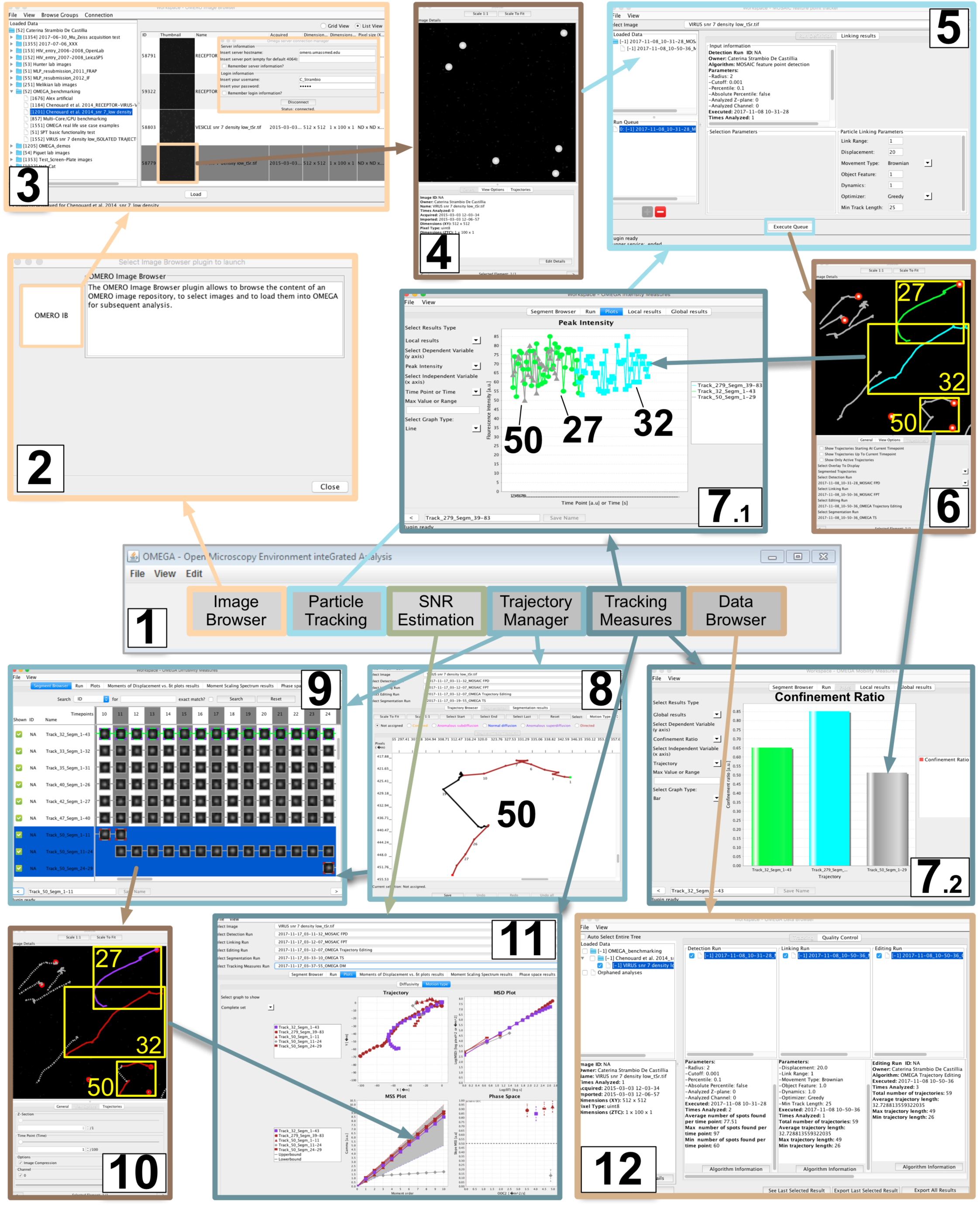
OMEGA for users: the graphical user interface (GUI). Upon starting the OMEGA application the user is presented with a tool-bar **(1)** offering access available plugins. After opening the Image Browser launch-pad **(2**) the user can launch the OMERO Image Browser plugin to select and load **(3)** one or more images of interest for inspection in the sidebar viewer **(4)**. After defining different paired Particle Detection and Particle Linking **(5)** runs the resulting spots and trajectories can be visualized as overlays via the sidebar image viewer **(6)**. Trajectories of interest (in the example trajectory nr. **27, 32** and **50**, as indicated here and in Figure 1) can be color coded (in the example trajectory nr. **27**, *green* and trajectory nr. **32**, *turquoise*) in order to facilitate their identification in all available views. After the execution of Intensity **(7.1)** and Mobility **(7.2)** Tracking Measures plugins, analysis results can be plotted using the same colors used on the sidebar to facilitate results comparison and interpretation. In case of non-uniform trajectories (*viz.* often observed with intracellular viral particles), they can be subdivided into individual segments of uniform mobility using the interactive user interface (GUI) provided as part of the Trajectory Segmentation plugin **(8)**. In the example, trajectory nr. **50** was subdivided in three segments: segments 1-11 and 24-29 were assigned a “directed” motion type (*maroon*), and segment 11-14, was left unassigned (*grey*). As a result of this color coding, when displayed on the Trajectory Browser **(9)**, the side bar **(10)**, and each of the Tracking Measures plot windows **(11)**, trajectory nr. **50** appears split in individual sections. Specifically, on the Trajectory Browser **(9)**, the initial color is used for the sides of the box enclosing each particle thumbnail (*grey,* in the highlighted example referring to trajectory nr. **50**); while the color corresponding to the motion type assignment is used for the vertexes of each box (*maroon*, in the highlighted example). In addition in order to facilitate results visualization, after segmentation trajectory segments whose motion type has not been yet assigned are displayed on the sidebar as dashed lines thus making it clear that they all belong to a single trajectory. At each step of the particle tracking and motion analysis workflow the user can maintain a clear picture of all available results stemming from different image-analysis data-flows using the OMEGA Data Brower **(12).** An additional advantage of this plugin is that it can be used to compare results from different runs as well as load results obtained from previous OMEGA sessions for further investigation.

One key feature of OMEGA that distinguishes from other tools (Table I) (Tinevez et al., 2016; de Chaumont et al., 2012). is that it provides experimental biologists wishing to quantitate in real time the movement of intracellular particles (i.e. vesicles, virions and organelles), with a unified data management tool capable of automatically keeping track and managing the complete data and provenance metadata flow from images to analysis output. Thus in addition to providing a strong interface with the OMERO image data and metadata repository (Allan et al., 2012), OMEGA also provides the infrastructure for automated data provenance harvesting, data-flow integration, standardized metadata annotation, uncertainty monitoring, and persistence of the whole analysis chain to facilitate dissemination, re-interpretation and meta-analysis of distributed particle tracking data. Furthermore, to facilitate reproducibility and results comparison across laboratories OMEGA carries out these functions within the framework of our newly proposed Minimum Information About Particle Tracking Experiments (MIAPTE) guidelines (Rigano and Strambio-De-Castillia, 2016; 2017), so that management, annotation, storage, and dissemination of the entire data cascade, is accomplished in a standardized manner. By unifying the entire image processing and analysis workflow, and by combining it with standardized data management and error propagation handling, OMEGA extends what is currently available, further reduces the need for users to transfer data manually across several downstream analysis tools, and lays the foundation for a particle tracking data dissemination and meta-analysis ecosystem.

All OMEGA algorithmic components were tested on artificial image and trajectory data as described in Supplemental Information 1 and elsewhere (Rigano et al., 2018). As proof of concept, OMEGA supported motion analysis workflows were utilized to analyzed both standard SPT benchmarking datasets (Figure 7; Chenouard *et al*, 2014), as well as real-life datasets depicting retroviral particle trafficking within living human cells (Clark et al., 2013; Pereira et al., 2012).

## 2 Tool description and functionality

### 2.1 Particle tracking and motion analysis workflow

In a typical particle tracking experiment (Figure 1A), time series of image-frames are recorded from living cells. If the image quality is sufficient, and the spatiotemporal resolution is adequate, SPT algorithms are then used to convert movies depicting particle motion (i.e., viral particles in the example shown) into statistical ensembles of individual trajectories specifying the coordinates and fluorescence intensity of each particle across time (Saxton, 2008; Rust et al., 2011). Subsequently, in a process that is often referred to as motion analysis (reviewed in, Meijering et al., 2012; Brandenburg and Zhuang, 2007), trajectories are used as input to compute quantitative measures describing the motion state of individual particles as well as their displacement, velocity, and intensity. The aim of this process is to correlate biochemical composition and functional readouts with particle dynamics and ultimately provide readouts that can be used to interpret the mechanisms governing motion and underlying intracellular interactions.

The OMEGA application (Figure 1; Supplemental Information 1) carries out all steps of the complete tracking and motion analysis workflow described above. OMEGA operates by way of a rich GUI (Figure 2; Supplemental Information 1), a modular structure, an automatic data provenance harvesting method and a robust persistence mechanism that relies on the OME-XML and the MIAPTE-compatible OMEGA data models to store data and metadata arising from image analysis either on the File System (FS) or on a dedicated relational database (Figure 3 and Supplemental Figure 2). The functional logic of the OMEGA Java application (Rigano and Strambio-De-Castillia, 2018a; Supplemental Information 1) is organized around six analysis and data management modular plugin-Superclasses (i.e., Image Browser, Particle Tracking, Signal-to-Noise Ratio -- SNR -- Estimation, Trajectory Manager, Tracking Measures, and Data Browser; Figure 3 and Supplemental Figure 2, solid lines boxes), which in turn are extended by twelve interchangeable plugins (Figure 3, dashed lines boxes) that work sequentially to execute the typical steps of particle tracking and motion analysis experiments, as well as to maximize results reproducibility and dissemination through the quantitation of motion type estimation uncertainty, and the recording of data provenance (Figures 1 and 4). At time of writing, workflows supported by OMEGA are mainly interactive, requiring user supervision at each subsequent step. In subsequent releases, we plan to develop batch processing of entire image datasets. Extensive validation of OMEGA components was conducted as described either here (Supplemental Information 1) or elsewhere (Rigano et al., 2018). Related error estimation and data provenance aspects are presented in parallel manuscripts (Rigano et al., 2018; Rigano and Strambio-De-Castillia, 2017).

**Figure 3:**
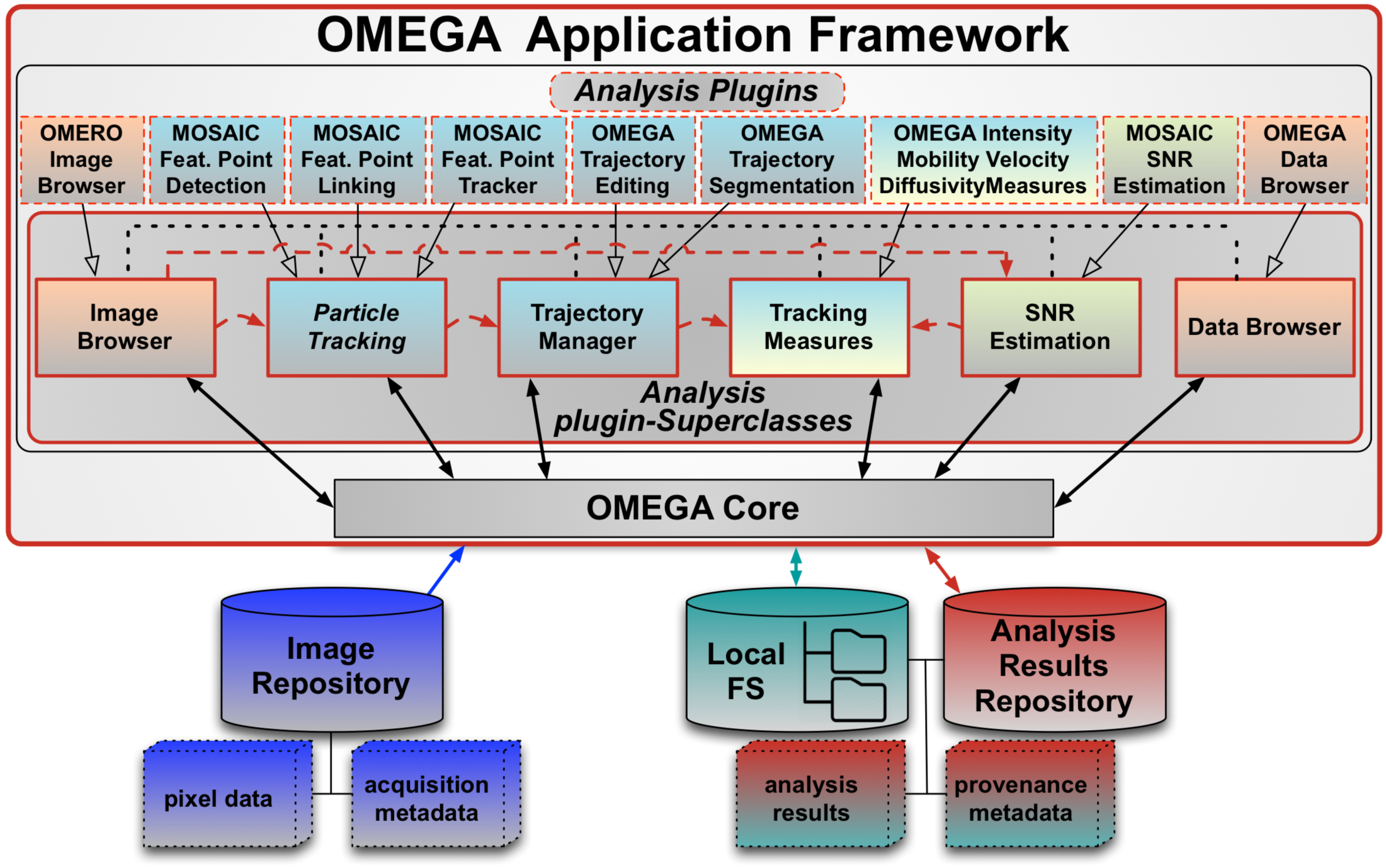
OMEGA’s modular software architecture. The OMEGA application framework is organized around the *OMEGA Core*, which imports image data and metadata from an associated *Image Repository* (*blue elements*), drives all processes, including those carried out by the *Analysis Plugins, and* writes output analysis results and provenance metadata (*red/green elements*) either to the file system (*Local FS; green cylinder*) or to the *Analysis Results Repository* (*red cylinder*). *OMEGA Core (*see also Supplemental Figure 2) contains all main sub-components of the application including the event-driven logic handling all communication between it and the *Analysis plugin-Superclasses* components. *OMEGA Core* also contains all GUI sub-components (Figure 2), including the plugin launcher dialog, which is opened every time users click on a button on the top toolbar. *Analysis plugin-Superclasses* contains all main functional logic in OMEGA. At the same time this component manages the flow of data between analysis plugins, automatically collects data provenance, and uses it to annotate output data (*dashed black line*). The *Analysis plugin-Superclasses* component is organized around six modular super-classes (solid borders boxes): 1) Image Browser; 2) Particle Tracking; 3) Trajectory Manager: 4) Tracking Measures; 5) SNR Estimation; and 6) Data Browser. Each of these modular types is extended (*empty black arrows*) by one or more interchangeable plugins, which in turn are responsible for specific steps of the analysis pipeline. At the time of writing, OMEGA ships with a set of nine Analysis (*cyan boxes*), two Data Management (*orange boxes*), and one dedicated Quality Control plugin (*light green boxes*). The Analysis plugins are: 1) *MOSAICsuite* Feature Point Detection (FPD; *MOSAIC Feat. Point Detection*); 2) *MOSAICsuite* Feature Point Linking (FPL; *MOSAIC Feat. Point Linking*); 3) *MOSAICsuite* Feature Point Tracker (FPT; *MOSAIC Feat. Point Tracker*); 4) OMEGA Trajectory Editing; 5) OMEGA Trajectory Segmentation; and 6-9) OMEGA Intensity-, Mobility-, Velocity- and Diffusivity-Tracking Measures. The Data Management plugins are: 1) OMERO Image browser; and 2) OMEGA Data browser. Quality control and error estimation are carried out by both the dedicated *MOSAICsuite* SNR Estimation plugin as well as by the motion-type estimation uncertainty components of the OMEGA Diffusivity Tracking Measures plugin (indicated by combined *cyan and light green shading*).

**Figure 4:**
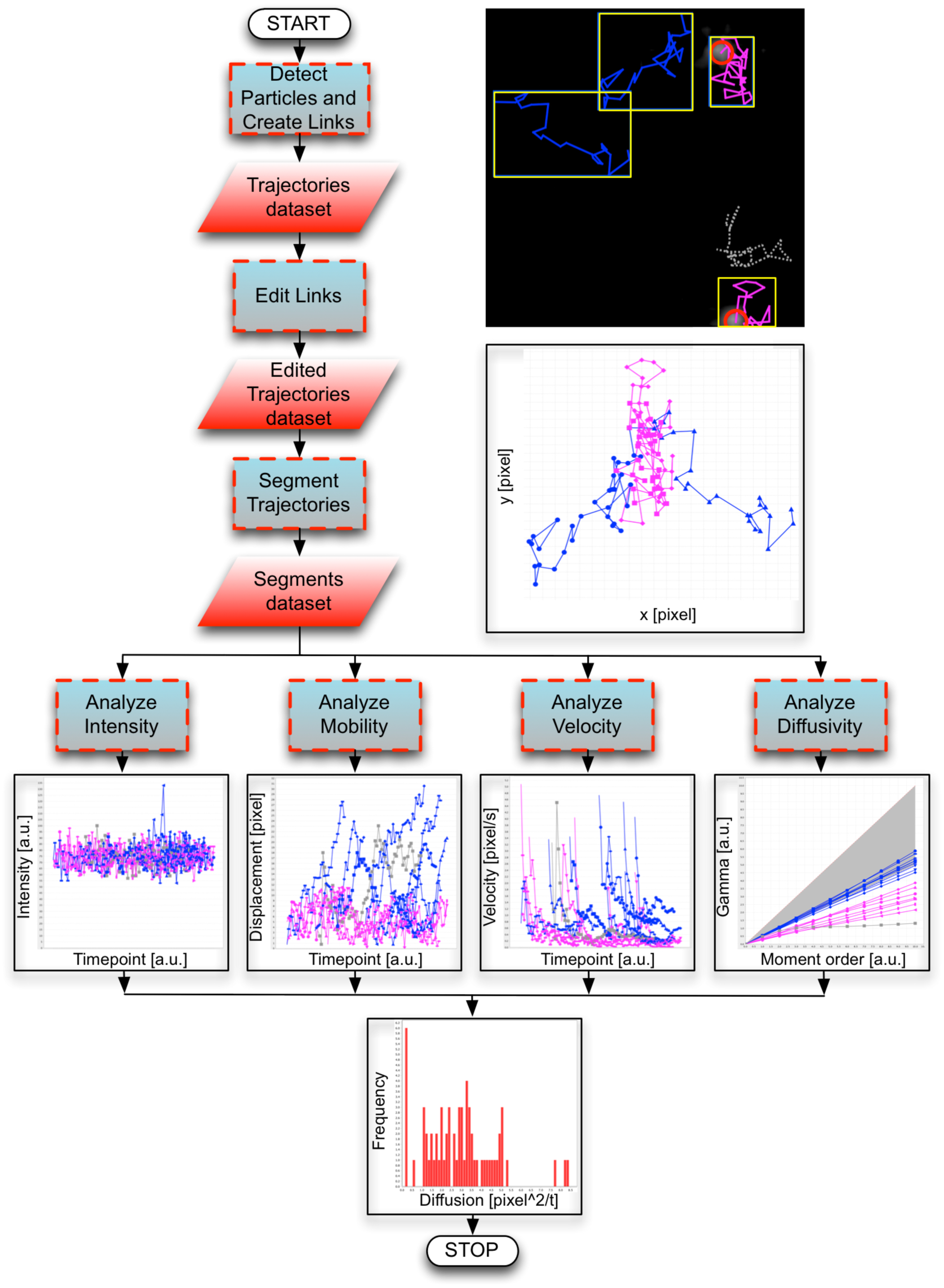
Motion analysis workflow in OMEGA. A flow-chart is employed to schematically depict the motion analysis workflow implemented in OMEGA. After particle detection and linking, when appropriate resulting trajectories datasets are subjected to link-editing and trajectory segmentation to produce validated and uniform trajectory segments. Segments are then analyzed using one or more of the available OMEGA Intensity Tracking Measures (ITM), Mobility Tracking Measures (MTM), VTM and DTM plugins. Finally, frequency distributions can be computed and data provenance-rich analysis results can be exported for more extensive statistical analysis using third party applications.

### 2.2 Data import

OMEGA supports two data import modalities, which are designed to assist users in the task of preserving data provenance links between different data processing steps. The first modality utilizes an Image Browser plugin (Figure 2-2 and 3) to import of image data and metadata from available repositories. The second modality relies on the OMEGA Data Browser (Figure 2-14 and 3; see below) to import and manage particle tracking and motion analysis data and metadata previously computed using OMEGA or other third party applications.

#### 2.1.1 Image Browser: importing images for particle tracking

At the time of writing, OMEGA is designed to import *BioFormats* (OME Consortium, 2017) compatible image data and metadata from the OMERO repository (Allan et al., 2012), via the custom-designed OMERO Image Browser plug-in (Figure 2-2 and 3). This plug-in provides a minimal interface to navigate through the OME Project, Dataset, and Image hierarchy, display available content in either a list or grid mode, select images to be analyzed and import them into OMEGA. This interface is specifically designed to visualize relevant image information while at the same time avoiding undue duplication of functionalities already available elsewhere. Only image thumbnails and metadata essential for image selection are shown, while additional viewing options are delegated to available OMERO clients (Allan et al., 2012). Plugins to import images from other data stores will be developed upon request.

#### 2.1.2 Data Browser: importing particle tracking analysis results

Among OMEGA’s centerpieces are built-in mechanisms to promote the reutilization of previously computed analysis results and interoperability. These mechanisms include the Data Browser (Figure 2-12 and 3; see below) import into OMEGA of MIAPTE-compliant (Rigano and Strambio-De-Castillia, 2016; 2017) particle tracking data and metadata computed beforehand either during previous OMEGA sessions or using third-party applications. Data can be imported either from the dedicated OMEGA Analysis Results repository or from the FS (Figure 3 and Supplemental Figure 2), and made available to the user via the Data Browser for all subsequent data navigation, selection and processing operations. If images corresponding to selected data are available in OMEGA, the imported data can be associated with them in the Data Browser. Alternatively, externally computed tracking data can be associated with a specifically designed “Orphaned Analyses” element. Examples of analyses that were performed on imported, pre-computed trajectories are presented in Figures 5 and 6B and Supplemental Figure 3.

**Figure 5.**
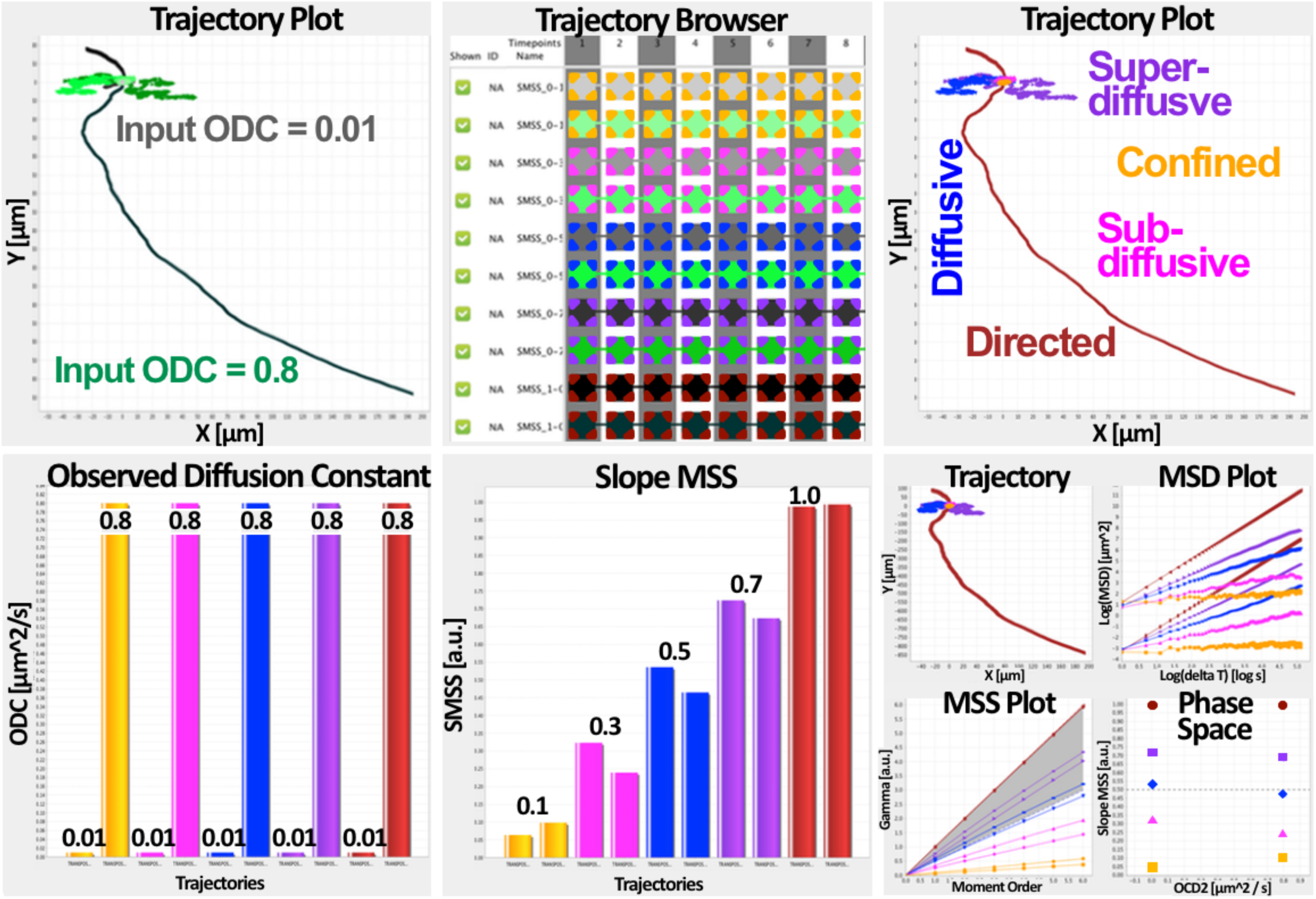
Motion type classification in OMEGA. Classification example using uniform artificial trajectories of known mobility characteristics. Ten self-similar artificial trajectories were generated using our *TrajectoryGenerator* MatLab algorithm (Supplemental Information 1). After importing into OMEGA (top left) using the Data Browser data importer, they were first arbitrarily color coded (i.e. shades of grey and green) and then assigned the motion type label corresponding to each expected motion type by using the Trajectory Segmentation plugin (top middle and right). Finally they were subjected to motion analysis using the DTM plugin (bottom row). Observed ODC (bottom left and right) and SMSS (bottom middle and right) numerical quantities and plot shapes were in excelled agreement with the corresponding indicated expected values. The position of each trajectory on phase space was consistent with the expected mobility (bottom right). Input ODC values as indicated: 0.01 and 0.8 Input SMSS values as indicated: 0.1, 0.3, 0.5, 0.7 and 1.0. Motion types color codes: *yellow*, confined; fuchsia, sub-diffusive; *blue*, diffusive; *purple*, super-diffusive; *maroon*, directed (Supplemental Table II).

**Figure 6.**
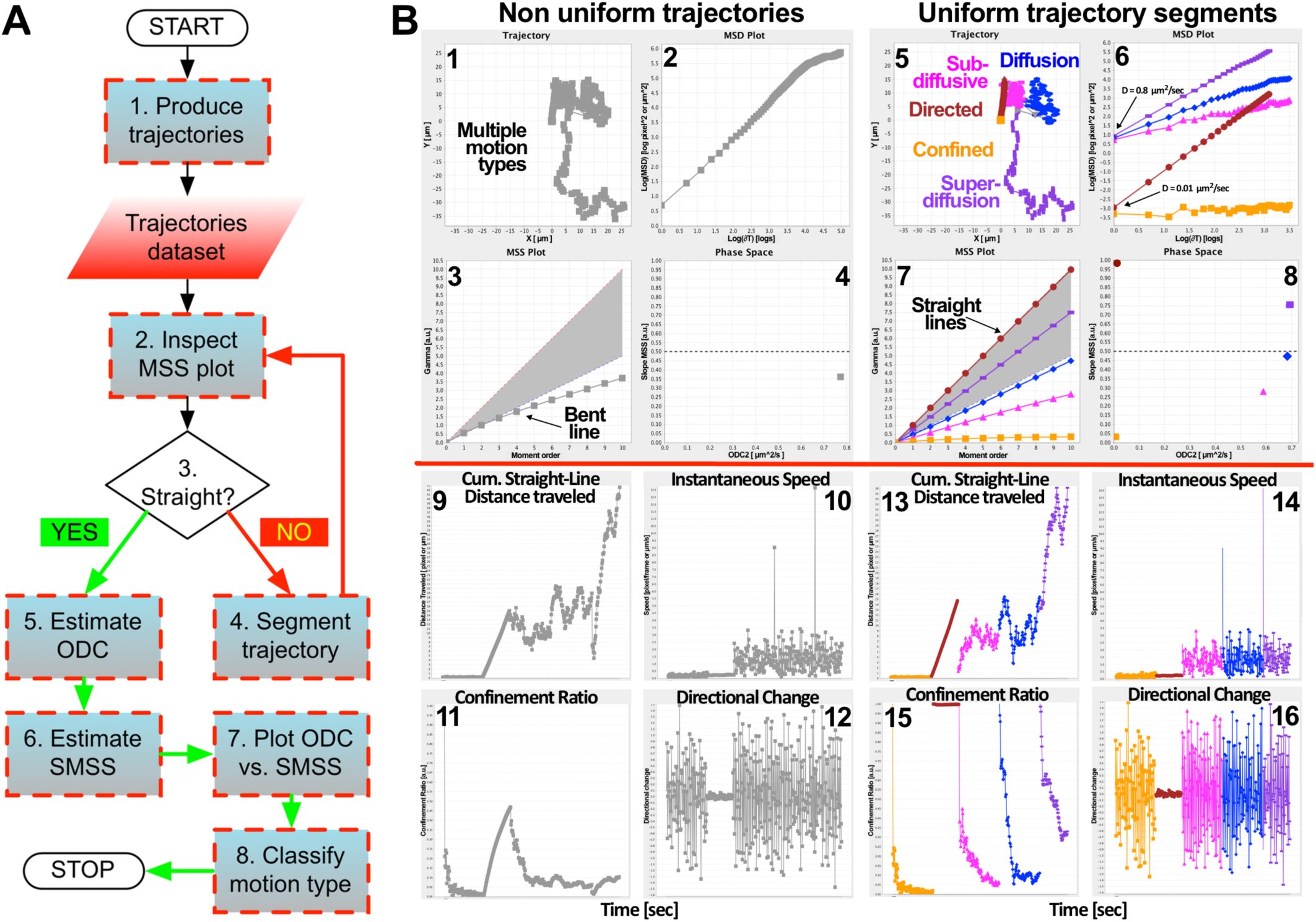
Iterative segmentation and classification workflow. A) Flow-chart diagram schematically depicting the iterative motion type classification workflow implemented in OMEGA. Step 1) The workflow starts with the production of trajectories by particle detection and linking. Subsequently, for each trajectory of interest the workflow continues as follows: Step 2) MSS analysis (Ferrari et al., 2001) and inspection of each MSS plot. Step 3) In case plots are bent, trajectories are subjected to Step 4, Else, the workflow continues to Step 5. Step 4) Trajectories are segmented and the process is repeated from Step 2. Step 5) ODC is estimated. Step 6) SMSS is estimated. Step 7) Each trajectory is plotted on ODC vs. SMSS phase space following the method of Ewers et al., 2005. Step 8) The motion behavior of each trajectory is classified based on where it falls on phase space. B) <u>Top left</u>: In order to validate the workflow presented in A, five uniform artificial trajectories of known ODC and SMSS as described in Figure 5 were merged to produce a single trajectory of mixed mobility (Non uniform trajectories). When subjected to the Ewers motion type classification method (Ewers et al., 2005) as implemented in the OMEGA DTM plugin (1 - 4), this mixed motion type trajectory gave rise to a clearly “bent” MSS curve indicating the non-uniform nature of the process (3). In this context, the calculated values of both ODC (2 and 4) and SMSS (3 and 4) represent averages over the length of the trajectory and are therefore not reliable. <u>Top right</u>: In order to correctly classify the behavior of each process component, the trajectory was subdivided in segments using the OMEGA Trajectory Segmentation plugin and each segment was analyzed individually (Uniform trajectory segments). As can be clearly observed (5 - 8), this gave rise to five independent straight MSS lines (7) indicating that the trajectory had been correctly subdivided in uniform segments, and allowing each segment to be analyzed independently from its neighbors (8). Bottom: Trajectories were subjected to motion analysis using the MTM (9, 11 and 12) and VTM (10) plugins, before (Non uniform trajectories) and after (Uniform trajectory segments) segmentation. When Straight-line Distance Travelled, Straight-line Speed, Confinement Ratio and Directional Change were plotted along each trajectory as a function of time, the resulting graphs reflected the presence of different motion components along the length of the full trajectory, which were correctly segregated after subdivision into individual segments.

### 2.3 Particle Tracking: detection and linking of spots to form trajectories

Once image time-series have been selected and uploaded into OMEGA, plugins extending the Particle Tracking super-class (Figure 2-5) convert movies depicting particle motion (Figure 2-4) into trajectories (Figure 2-6) consisting of the coordinates and fluorescence intensities of each particle across time (Saxton, 2008; Rust et al., 2011). Particle Tracking is generally subdivided into two independent steps: 1) Particle Detection identifies individual fluorescent spots that are significantly distinguishable from local background and estimates their sub-pixel coordinates and intensities in each time frame. 2) Particle Linking generates trajectories by linking the position of each bright spot in one time frame with its position in subsequent frames.

Recent systematic and objective comparisons of available particle tracking algorithms on standardized benchmarking data sets (Chenouard et al., 2014; Saxton, 2014), have shown that no single algorithm or set of algorithms can optimally solve all tracking problem sets. What is clear is that for any given experimental context and scientific question, multiple algorithms or combinations thereof should be rigorously evaluated before moving to production. This situation makes parameterization and testing of algorithms an essential component of the process. To facilitate this task the OMEGA Particle Tracking super-class is designed to accommodate three different plugin designs: 1) Particle Detection stand-alone; 2) Particle Linking stand-alone; and 3) integrated Particle Tracking plugin merging both detection and linking functionalities into a single element. This flexible plugin strategy facilitates the integration of diverse tracking algorithms regardless of their specific implementation style and allows the user to “mix and match” compatible particle detection and linking algorithms originating from different sources with a significant improvement in tracking quality and efficiency. In addition, the architecture of the Particle Tracking super-class in OMEGA emphasizes modularity and open APIs in order to facilitate the integration of different third-party detection and linking algorithms to be tested on each experimental case. Last but not least, the Data Browser component of OMEGA (Figure 2-14; see below) allows the user to store not only the results of different SPT runs, but also all associated parameter-settings metadata. This enormously facilitates the systematic comparison of runs and the selection of the best tracking method for a given experimental situation of interest.

As a proof of concept, we ported into OMEGA the *MOSAICsuite* Particle Tracker ImageJ plugin (Sbalzarini and Koumoutsakos, 2005; Incardona and Sbalzarini, 2014). This algorithm was selected because of its versatility and computational efficiency as formally assessed in a recent side-by-side comparison using benchmarking datasets (Chenouard et al., 2014). In addition, the algorithm was specifically designed for images with low SNR where prior knowledge of motion modalities is absent, such as during the exploration and optimization phases of a tracking assay. Because the *MOSAICsuite* Feature Point Tracker (FPT; Incardona and Sbalzarini, 2014) carries out both particle detection and linking, in addition to creating an equivalent dual-function plugin for OMEGA, the underlying algorithm was split to generate two stand-alone *MOSAICsuite* Feature Point Detection (FPD) and *MOSAICsuite* Feature Point Linking plugins (FPL; for validation details see Supplemental Information 1). The integration of additional Particle Detection, Particle Linking, and Particle Tracking plugins in OMEGA is planned for future releases.

### 2.4 SNR Estimation: image quality control

The accuracy and precision of particle detection as well as that of all downstream trajectory analysis steps depend very closely on the local SNR observed in the immediate surroundings of each identified particle. The OMEGA SNR Estimation module is extended by plugins that estimate the SNR associated with each tracked particle allowing the identification of images whose quality does not support reliable particle detection and tracking. At time of writing, OMEGA ships with a single local SNR Estimation plugin consisting of a custom Java implementation of the *MOSAICsuite*’s local SNR Estimation algorithm originally developed by the MOSAIC group in Matlab (Xiao et al., 2016; Gong and Sbalzarini, 2016; Rigano et al., 2018; Rigano and Strambio-De-Castillia, 2018a). This routine, which was benchmarked as described (Supplemental Information 1), uses the particle coordinates obtained from the Particle Detection plugin to extract intensity values from each associated image plane and estimate background, noise and SNR pertaining to the area immediately surrounding each particle. Moreover, the plugin computes aggregate SNR values at both the plane and the image level. Briefly, the algorithm first determines the global background and noise associated with the entire image plane where individual particles are localized. It then takes the particle radius as defined by the user to draw a square area around each particle’s centroid and identify the brightest pixel within the particle area (i.e., peak intensity). Finally, it estimates local noise, local background and local SNR. Specifically the latter is calculated using three independent models: two are based on the Bhattacharyya distance (i.e., Bhattacharyya Poisson, and Bhattacharyya Gaussian) and the third is based on the method proposed by Cheezum (Cheezum et al., 2001).

After calculating local SNR values, the algorithm returns aggregate SNR values at the trajectory, plane and image level, which are utilized two-fold in OMEGA. The global local SNR average over the entire image is used to estimate the minimum detectable Observed Diffusion Constant (ODC; Supplemental Information 1). The global minimum local SNR each given trajectory is utilized to estimate the confidence associated with both ODC and SMSS estimations and with motion type classification as described (Rigano *et al*, 2018). In addition, image averages, minimum and maximum local SNR values can be utilized by the user to evaluate the overall image quality and general performance of the particle detector as specified by the algorithm developer (see below). For example, in the case of the *MOSAICsuite* Particle Detection algorithm, which is currently implemented in OMEGA, an SNR value lower than a threshold of 2 as calculated according to Cheezum (Cheezum et al., 2001), indicates to the user that particle detection and motion analysis will not be reliable and better images should be acquired.

### 2.5 Trajectory Manager: trajectory curation

This super-class, which is extended by Trajectory Editing and Trajectory Segmentation plugins, provides interactive graphical support for the inspection of trajectory data quality, correction of linking errors and subdivision of trajectories in segments of uniform mobility.

#### 2.5.1 Trajectory editing

Linking algorithms generally perform satisfactorily provided certain SNR, spatiotemporal sampling and observation times criteria are met (Jaqaman and Danuser, 2009). However, despite steady improvements in particle linking methods (Sbalzarini and Koumoutsakos, 2005; Chenouard et al., 2014; b; c; Jaqaman et al., 2008; Sergé et al., 2008; Tinevez et al., 2016; Genovesio et al., 2006; Ku et al., 2007; Jug et al., 2014; Kalaidzidis, 2009) it is often necessary to manually verify individual links. The most frequent linking errors are due to excessive particle density; insufficiently temporal sampling (i.e., particles move too fast with respect to time interval employed during acquisition); splitting or merging of trajectories, which might result either from an artifact (i.e., two or more particles are close enough that their distance is below the diffraction limit) or from the actual interaction between particles; and particle blinking or moving temporarily out of focus. The OMEGA Trajectory Editing plugin uses our custom Trajectory Browser graphical interface to facilitate splitting and merging of trajectories that upon inspection appear to be faulty (Figure 2-9).

#### 2.5.2 Trajectory segmentation

The movement of intracellular objects, such as viral particles or vesicles, is often characterized by frequent switches between different dynamic behaviors the analysis of which can be used to infer interactions between the moving object and its immediate surroundings (Helmuth et al., 2007). For example the interaction of viral particles with motor proteins might result in directed motion along microtubules (Brandenburg and Zhuang, 2007; Arhel et al., 2006; McDonald et al., 2002; Sun et al., 2013; Fernandez et al., 2015). On the opposite side of the spectrum, interaction with relatively immobile cellular structures such as nuclear pore complexes or membrane rafts, might result in the transient confinement of particles to restricted zones (Schelhaas et al., 2008; Burckhardt and Greber, 2009; Kusumi et al., 2014; 1993). Because motion characteristics can be reliably estimated only when trajectories describe stationary and ergodic processes, it is often necessary to decompose trajectories into individual uniform polyline segments to be individually subjected to motion analysis. One additional advantage of this process, herein referred to as trajectory segmentation is that events can be defined as specific series of segment types whose frequency can chance as a result of specific molecular or cellular events.

The Trajectory Segmentation plugin of OMEGA provides a specialized version of the Trajectory Browser GUI to assists the user in decomposing trajectories into uniform tracts, each of which can then be analyzed separately (Figure 2-8). The tool allows users to select manually the start and end point of segments and assign a putative motion type to each segment (i.e., *yellow*, confined; fuchsia, sub-diffusive; *blue*, diffusive; *purple*, super-diffusive; *maroon*, directed; Supplemental Table II; Figures 5 and 6B; Supplemental Figure 3). This manual method can be used in conjunction with an iterative analysis process to obtain homogeneous trajectory segments (see below and Figure 6). In subsequent releases, automated methods for trajectory segmentation will be integrated in OMEGA (Helmuth et al., 2007; Wagner et al., 2017; Huet et al., 2006; Persson et al., 2013; Wang et al., 2017).

### 2.6 Tracking Measures: trajectory analysis, motion type classification and error estimation

Trajectory analysis reduces a sequence of spatial coordinates into scalar quantification parameters that are computed using various averaging techniques applied along the length of the trajectory (Supplemental Table I). The ultimate goal is to gain new understanding about the system under study by computing “biologically meaningful quantitative measures from these coordinates” (verbatim from: Meijering et al., 2012). Specifically, OMEGA computes Intensity Tracking Measures (ITM), Mobility Tracking Measures (MTM) and Velocity Tracking Measures (VTM), which are only subject to localization accuracy (i.e., algorithmic systematic bias) and precision (i.e., noise associated random errors) and are therefore considered deterministic (Figures 1, 2 and 4). In addition, OMEGA calculates quantities such as Diffusivity Tracking Measures (DTM) whose values are strongly influenced by sample size, and are therefore statistical (Figures 1, 2 and 4). To facilitate all analysis tasks, the OMEGA Tracking Measure plugins provide a rich interface for users to select trajectory segments using a specialized Segment Browser panel (Figure 2-9), perform quantitative motion analysis on selected segments, examine results in both a tabular and graphical form, export data for downstream processing using third-party application and produce publication grade figures (Figures 4-8). All Tracking Measures plugins (Figure 4) calculate and display both local (i.e., pertaining to an individual particle) and global measures (i.e., pertaining to a whole trajectory or trajectory-segment) as applicable. Furthermore, these plugins can perform limited statistical analysis by computing frequency distributions at the image level. In order to compare results obtained across different images and datasets the user can export results and perform downstream statistical analysis using tools such as R or Matlab (MATLABMathWorks:2018tc; The R Foundation, 2018).

In addition to calculating tracking measures, OMEGA facilitates the classification of either full trajectories or individual uniform segments based on motion type using the ODC vs. SMSS phase space method developed by Ewers et al. (Figures 5 and 6; Supplemental Figure 3; Supplemental Table II; Ewers et al., 2005; Schelhaas et al., 2008; Sbalzarini and Koumoutsakos, 2005). Last but not least, OMEGA estimates the effect of spot detection uncertainty, limited trajectory length and motion type on the reliability of downstream motion analysis results as an essential pre-requisite for scientists to critically evaluate and compare results (see below).

#### 2.6.1 Trajectory analysis

##### Intensity Tracking Measures

Fluorescently labeled particles might display changes in fluorescence intensity as a result of addition or loss of labeled components as well as of photo-bleaching and -toxicity. It is therefore often important to report changes in the signal intensity of tracked objects (Figure 4 and Supplemental Table I). For example, when tracking dual-color enveloped viral particles carrying a membrane marker, a sudden drop in intensity might indicate fusion between the viral envelope and the acidic endocytic compartment (Mamede et al., 2017; Sood et al., 2017; Padilla-Parra et al., 2013; Sood et al., 2016; Itano et al., 2018). Alternatively, when studying endocytic trafficking, changes in intensity might inform about specific cargo sorting events (Navaroli et al., 2012). Most tracking algorithms compute either centroid intensity, peak intensity or both. In addition, the mean intensity of the particle might also be computed when the area of the particle is available. The OMEGA ITM plugin gathers relevant intensity values for each identified particle either directly from the Particle Detection plugin or if necessary from the local SNR Estimation plugin and makes them available for visualization on screen as well as for downstream analysis.

##### Mobility Tracking Measures

Mobility measures are relatively easy to compute and assess the quantity of motion away from the origin or from a reference point, the duration of motion and the persistence along a specific direction (Figures 4 and 6; Supplemental Figure 3; Supplemental Table I). For example, in viral trafficking it is important to quantify what proportion of viral particles move consistently towards the cell center versus those that remain confined near the site of viral entry at the cellular periphery (Yamauchi et al., 2011; Navaroli et al., 2012; Jaqaman et al., 2016; 2011). In OMEGA, local MTM quantify motion associated with a single step (i.e., trajectory link) and include Distance Traveled, Instantaneous Angle and Directional Change. Among global measures computed in OMEGA are: Total Curvilinear Distance Traveled (i.e., the total path length followed by a Brownian particle from start to end of motion; (Saxton, 2009), the Total Net Straight-Line Distance Traveled (i.e., the distance between the beginning and the end of motion) and the Confinement Ratio (also known as the meandering index, straightness index or directionality ratio) as a measure of the straightness of trajectories or the confinement of moving particles (Sergé et al., 2008; Beltman et al., 2009).

##### Velocity Tracking Measures

Measuring the rate of displacement of moving intracellular objects such as vesicles or viral particles can provide important information about the underlying mechanism of motion (Figures 4 and 6; Supplemental Figure 3; Supplemental Table I). For example, a fast moving viral particle that is also moving in a consisted direction might be moving along microtubules while a particle confined within a membrane raft might remain relatively still for large proportions of time (Gazzola et al., 2009; Suomalainen et al., 2001; Engelke et al., 2011). In OMEGA, VTM include local measures such as Instantaneous Speed. Global measures include Average Curvilinear Speed, Average Straight-line Speed, and Forward Progression Linearity, which gives a measure of how quickly an object is moving away from its origin during the total trajectory time (Meijering et al., 2012).

##### Diffusivity Tracking Measures

Because of their size, intracellular vesicles, virions and all diffraction-limited objects behave like Brownian particles, whose default state is normal diffusion (Saxton, 2009). Under these conditions, alteration in the diffusion state of particles result from molecular interactions that alter the general direction or the rate of motion. As a corollary, the distinction between normal vs. abnormal diffusion represents one of the most important steps in an attempt to distinguish between phases in which particles are free to move about and phases in which they are engaged in interactions with the surrounding cellular milieu that restrict their mobility. This in turn provides important clues for the understanding of the underlying mechanisms influencing the behavior of intracellular virions, vesicles, and other structures.

All motion processes can be described in terms of the *probability* that a given particle that at time 0 is found at position x(0), moves to position x(t) at time t. Thus, despite the fact that the diffusion coefficient (D; Saxton and Jacobson, 1997) is not constant in time for anomalous diffusion processes, this allows the extension of diffusivity analysis to all types of anomalous diffusion (Ferrari et al., 2001; Sbalzarini and Koumoutsakos, 2005). Based on these premises, OMEGA implements a single method to classify the dynamic behavior of individual particles regardless of their motion characteristics and employs the same method for particles whose dynamic behavior changes during the course of motion, as is commonly observed in living systems (Tables I and II).

Specifically, the method implemented in OMEGA (Figures 4-8; Supplemental Figure 3; validated as described in Supplemental Information 1) reproduces well-known methods (Sbalzarini and Koumoutsakos, 2005; Schelhaas et al., 2008; Ewers et al., 2005), which combines two components: 1) quantitative assessment of the degree to which the motion characteristics of the particle under study deviate from free diffusion; and 2) estimation of the quantity of displacement. In order to assess the diffusivity characteristics of a given particle, the OMEGA DTM plugin (Figures 4-8; Supplemental Figure 3) uses a well-established method based on the observation that the Squared Displacement (SD) of a diffusing particle from the origin of motion grows linearly with time in expectation. After time averages of SD (Landau and Lifshitz, 1960) – Mean Squared Displacement (MSD) – are computed for individual trajectories as a function of calculation time lag (Δt), the scaling behavior (i.e., slope) in plots of log(MSD) *vs.* log(Δt) can be used to calculate D and is sometimes used as an indication of whether the trajectory under study is characterized by normal (slope = 1) or anomalous diffusion (slope ≠ 1; Supplemental Table II; Saxton, 1993).

Because the slope of log(MSD) *vs.* log(Δt) plots is not sufficient to discriminate between normal and abnormal diffusion, OMEGA implements a method developed by Ferrari et al. and based on the estimation of the Hurst exponent (Turner et al., 2009; Hurst, 1951; Ferrari et al., 2001) to increase the accuracy of motion type prediction. This method, primarily referred to as Moment Scaling Spectrum (MSS) analysis, extends the study of the logarithmic scaling behavior with respect to Δt to moments of displacement (μν) other than the moment of displacement of order ν = 2, which corresponds to the MSD (i.e., MSD = μ_2_). Thus, a MSS graph is constructed plotting the values of μν vs. the corresponding values of the logarithmic scaling factor (i.e., ϒν) and the Slope of the MSS curve (SMSS) is used to discriminate between different motion modalities, where SMSS = 0.5 corresponds to normal diffusion, while values of SMSS ≠ 0.5 correspond to anomalous diffusive states (Supplemental Table II). In order to calculate the quantity of displacement, OMEGA calculates the generalized Observed Diffusion Constant of order ν = 2 (ODC_2_), which in case of a purely diffusive Brownian particle coincides with D (Saxton and Jacobson, 1997).

Specifically, OMEGA calculates the values of μν for ten different orders (ν, as well as all corresponding ϒν and ODCν values, and reports them in tabular form. In addition, OMEGA reports values of ODC2 (i.e., henceforth referred to as ODC) calculated from the intercept of the linear regression of log-log (ODC2log) plots of MSD vs. Δt (Sbalzarini and Koumoutsakos, 2005), which is more robust in case of trajectories that differ significantly from free normal diffusion. Finally, the value of the slope of the log-log plot of MSD vs. Δt (ϒ2), as well as the SMSS (also termed β) are reported as global estimates of the dynamic behavior of particles under study and made available in both graph and table format (Figures 4-8; Supplemental Figure 3; Supplemental Table I).

#### 2.6.2. Motion type classification

Global motion analysis reduces whole trajectories to a series of individual measurements (Tables I and II). The combination of two or more of such features enables the representation of individual trajectories as points in n-dimensional phase space (Schelhaas et al., 2008; Ewers et al., 2005; Sbalzarini and Koumoutsakos, 2005). In addition to representing a massive data reduction, this approach has the advantage of facilitating the classification of the mobility characteristics of multiple particles all at once without arbitrary selection. Thus, trajectories clustering in phase space are expected to have similar dynamic behavior and in turn correspond to similar functional states. An additional advantage of this method is that states described in this manner could be defined dynamically depending on individual scientific questions while at the same time could be the subject of standardization. At the time of writing, motion classification in OMEGA is based on the phase space of SMSS *vs.* ODC, which allows to quantify both the “speed” and the “freedom” of a group of moving objects independently, as previously described (Schelhaas et al., 2008; Ewers et al., 2005; Sbalzarini and Koumoutsakos, 2005). In order to test this approach, uniform artificial trajectories of known mobility were generated (Supplemental Information 1) using the custom-made *TrajectoryGenerator* Matlab routine (Helmuth et al., 2007; Rigano and Strambio-De-Castillia, 2018b), imported into OMEGA using the OMEGA Data Browser plugin import feature, and assigned the motion type label corresponding to defined motion type classification criteria (Supplemental Table II) by using the OMEGA Trajectory Segmentation plugin. After subjecting to diffusivity analysis using the OMEGA DTM plugin, trajectories were plotted on phase space on the basis of their measured ODC and SMSS, which resulted in excellent agreement with the ground-truth behavior (Figure 5 and Supplemental Figure 3).

The addition of further features to augment phase-space clustering, such as the Directional Change between subsequent displacement steps or measures of anisotropy (Huet et al., 2006) is easily implementable due to the generic nature of the underlying architecture and is planned for future releases.

##### Iterative motion type classification/segmentation workflow

As mentioned, motion type classification in OMEGA is based on the visual inspection of log-log plots describing the variation of either MSD or of moments of displacement of different order over increasing time intervals (Supplemental Table II). In the presence of motion type transitions within an individual trajectory (e.g. periods of confinement followed by normal diffusion; or periods of deterministic drift interspersed with bursts of directed motion), global quantitative measures that are averaged over the entire duration of the trajectory, such as ODC and SMSS, represent unreliable estimates of particle dynamics (Ewers et al., 2005; Helmuth et al., 2007). To obviate this hurdle, OMEGA implements an interactive pipeline for motion type classification (Figure 6A), which is based on the notion that the MSS plot carries information about the “self-similarity” of the motion under study (Ferrari et al., 2001). Thus, if all moments in the spectrum scale linearly with order, then the MSS is a line and the motion is defined as “strongly self-similar”; conversely if the MSS curve is kinked or bent, the movement is classified as “weekly self-similar”, indicating the existence of transitions between different states. Specifically (Figure 6A): 1) after particle tracking, trajectories are subjected to MSS analysis. 2) If the resulting plot is observed to be bent, the trajectory can be iteratively subdivided into segments until all resulting segments produce a straight MSS line. 3) At this point, ODC and SMSS are estimated and each segment is plotted as a point in phase space. The position of each trajectory in phase space as described above reflects their dynamics and is used to assign trajectories to motion type classes whose frequency in the segment population can be estimated by drawing windows around clouds of points, and compared across experimental conditions by statistical analysis using third-party applications as a prerequisite for functional analysis.

In order to validate the iterative segmentation vs. classification workflow, uniform artificial trajectories (Supplemental Information 1; Rigano and Strambio-De-Castillia, 2018b; Helmuth et al., 2007) were merged to produce trajectories comprising five different motion types (Figure 6B). This resulted in “bent” MSS curves, indicating the non-uniform nature of the overall process. When the mixed trajectory was subdivided in segments and each was analyzed individually, this gave rise to five independently linear MSS curves allowing each segment to be correctly classified independent of its neighbors.

#### 2.6.3 Estimation of motion type classification error

In order to interpret and draw valid conclusions from analysis results, the error associated with each measurement or calculation has to be determined and its effect on downstream analysis steps (i.e., error propagation) has to be clearly understood. Despite the apparent truism of this statement, attention to error propagation in particle tracking has been limited (Sbalzarini, 2016). To address this issue, OMEGA incorporates both theoretical and empirical methods to estimate uncertainties associated with motion analysis results. Details of our error estimation procedure are described in an associated manuscript (Rigano et al., 2018), here we provide a short description of the method.

While linking errors are orthogonal to motion analysis and are addressed elsewhere (Tinevez et al., 2016), uncertainty associated with spot detection strongly affects the accuracy with which trajectories can be classified on the basis of their observed dynamic behavior (Ewers et al., 2005). In addition, even with infinitely precise and true positioning, trajectory measures are expected to display statistical uncertainty because of finite trajectory lengths. These finite-data uncertainties diminish as the number of points that are detected as part of each trajectory increases. In addition to these well-known sources of error (i.e., position and sampling), we also determined that the quantity of displacement as well as the freedom of motion of moving particles (i.e., ODC and SMSS respectively) affect motion type estimation uncertainty.

When image quality is low (i.e., low SNR), the verisimilitude of positioning estimates might be as low as to make it difficult to distinguish between actual movement and apparent positional shifts arising from both systematic and random localization errors. This is particularly problematic for random walks, which can be discriminated from sub-diffusing particles and even from stationary particles only when their ODC is large enough to cause particle motion larger than the localization uncertainly (Martin et al., 2002). Based on these premises, given the observed image quality it is possible to define a “limit of detection” below which ODC values can be considered meaningless. For this purpose we employ the global error model described by Martin et al. (Martin et al., 2002) to calculate Minimum Detectable ODC of order 2 (*ODC*_2*MinDet*_) values as a function of image quality and detection (see Supplemental Information 1). Once calculated, *ODC*_2*MinDet*_ is reported in both tabular and graphical form (Figures 7 and 8). This threshold can be used to exclude from subsequent analyses steps trajectories whose global ODC value is too low to be meaningfully distinguished from noise.

**Figure 7.**
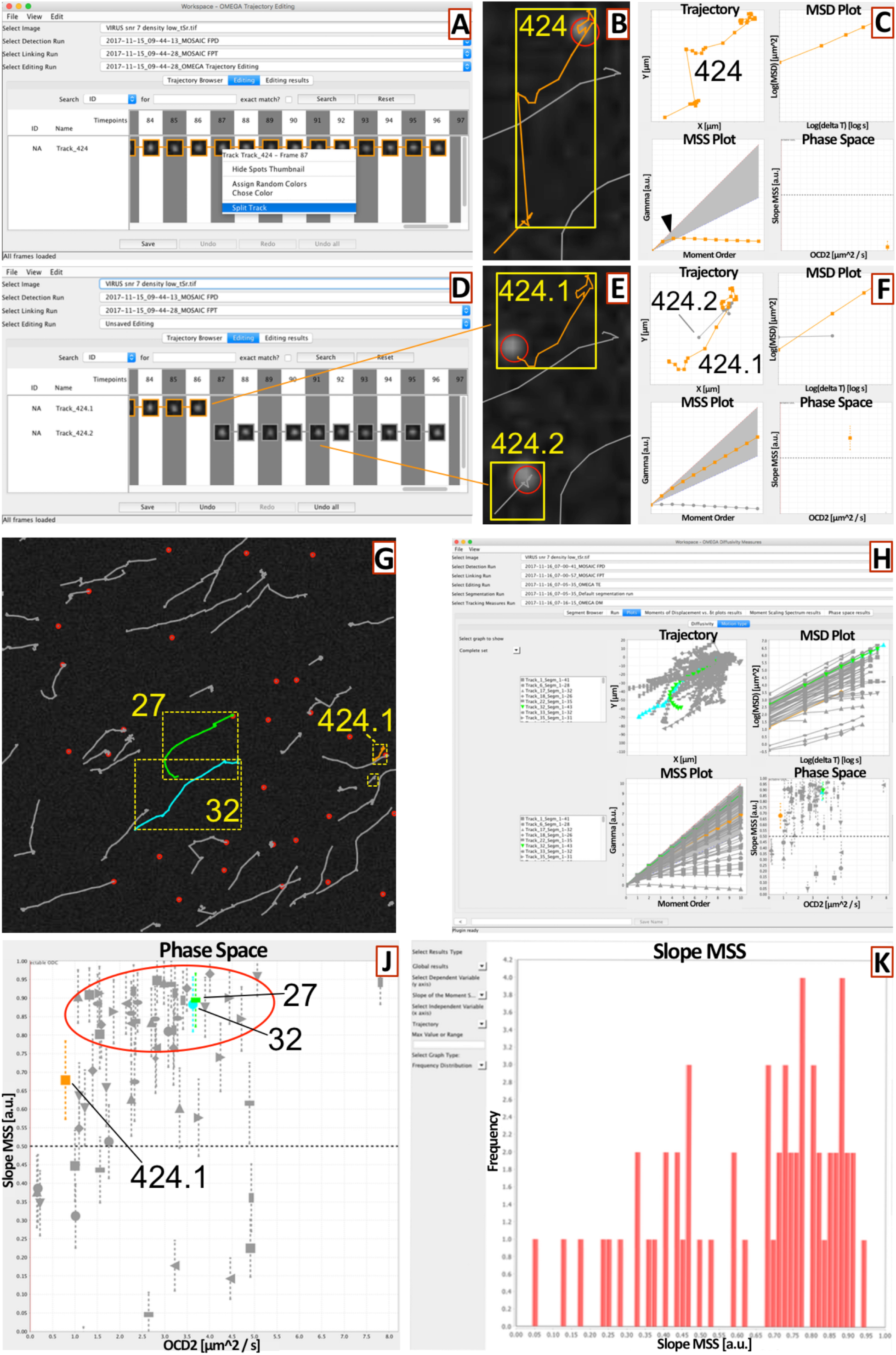
Example use-case using standardized SPT benchmarking datasets mimicking viral particle movement in infected cells. Time series image from the Chenouard et al. SPT benchmarking dataset (Chenouard *et al.*, 2014) corresponding to scenario IV, SNR = 7 and low particle density, was subjected to SPT within OMEGA. As expected most resulting trajectories displayed a behavior mimicking Levy flights (Levy, 1937), such as what observed in active motion. While most trajectories appeared to be valid, trajectory nr. 424 appeared to be the result of two erroneously linked particles, which was also confirmed by the appearance of a clear bend in the MSS curve (panel C, bottom left, *arrowhead*). When trajectory nr. 424 was edited using the Trajectory Editing plugin (panels A and D), the resulting allowed to split this trajectory in two individual trajectories 424.1 and 424.2 (panels E and F), one of which (nr. 424.1) gave rise to a straight MSS curve, consistent with the behavior of a uniformly mobile particle (panel F). After editing, trajectories were subjected to diffusivity analysis (panels G-K), yielding trajectories clustering in the top quadrant of the phase space graph (panel J, *red circled area*) as well as the prevalence of SMSS values close to 1 (panel K).

**Figure 8.**
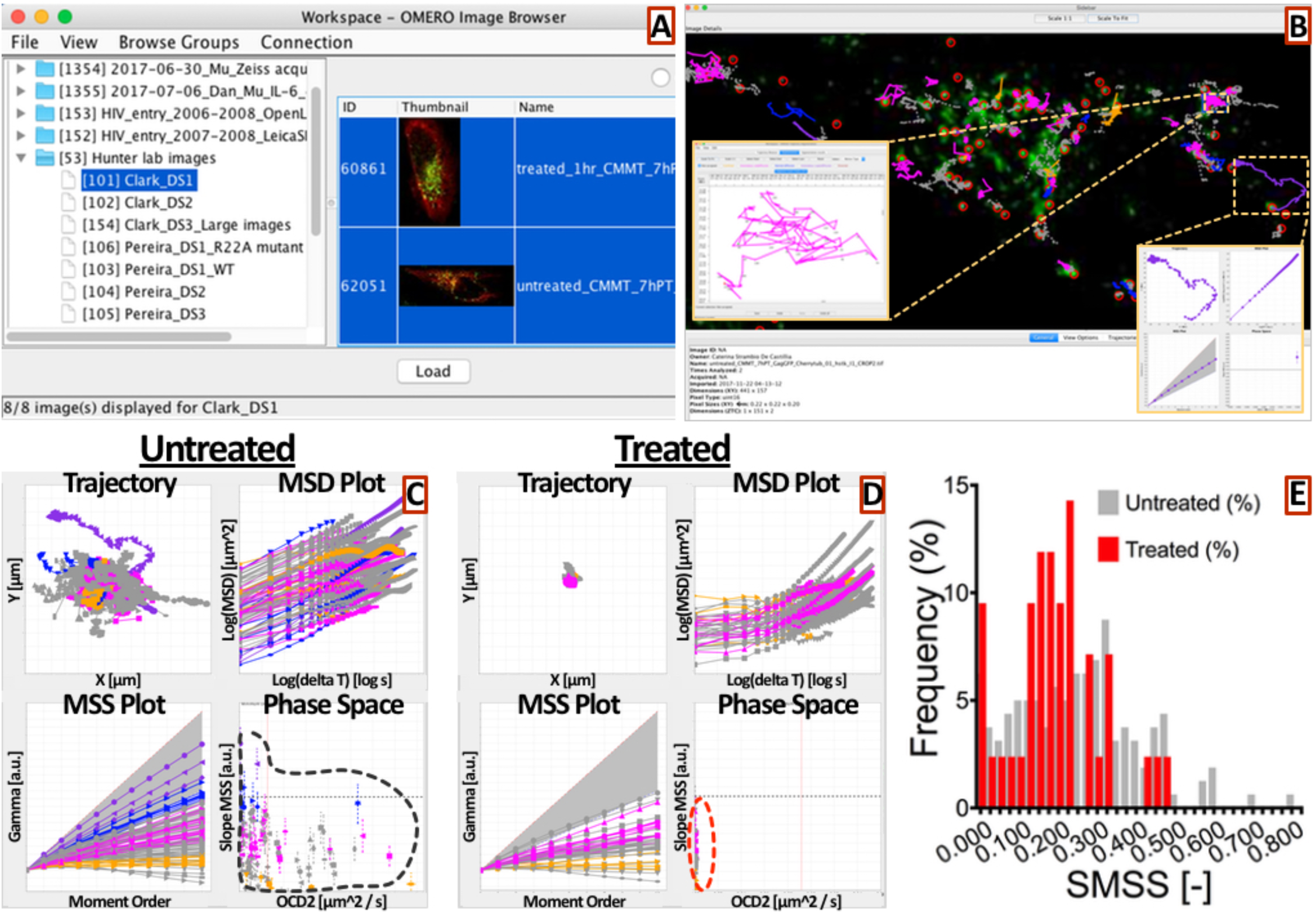
Example use-case using real-life imaging data: treatment with Nocodazole drastically reduces Gag-containing viral particles during M-PMV viral assembly. M-PMV producing, rhesus macaques CMMT cells were co-transfected with pSARM-GagGFP-M100A and p-mCherry-Tubulin to visualize the cytoplasmic assembly and trafficking of immature viral particles (Clark *et al.*, 2013; Pereira *et al.*, 2012; Pereira *et al.*, 2014). 7 hr post-transfection cells were either mock-treated (*Untreated*) or treated with Nocodazole for 1 hour (*Treated*), before microscopic observation under 60X magnification using a Delta vision deconvolution fluorescence microscope (Applied Precision Inc., Issaquah, WA), equipped with a Cool Snap CCD camera. All acquisitions were performed at 37°C in a micro chamber with CO2 infusion. 3D images (with 10 z-focal sections spaced 200 nm apart) were in captured every 5 seconds for a total of 2 minutes. Images presented here are maximum projections of all z-sections in one plane. (A) After acquisition images were loaded onto OMERO and imported into OMEGA using the OMEGA image browser. (B) Images were subjected to SPT using the OMEGA implementations of the *MOSAICsuite FPD and FPL* plugins (Sbalzarini & Koumoutsakos, 2005). The resulting particles and trajectories were overlaid over the corresponding image using the OMEGA side bar image viewer. All trajectories were subjected to diffusivity analysis using the OMEGA DTM plugin. Trajectories that displayed a straight MSS plot curve were assigned the corresponding motion type using the OMEGA Trajectory Segmentation plugin. Insets display the motion-type assignment graphical user interface for a representative sub-diffusive (*fuchsia*) viral particle and the motion type classification 4-plots set (xy, MSD vs. t log-log, MSS and D vs. SMSS phase space plots) for a representative super-diffusive (*purple*) trajectory. (C) Global view of all identified trajectories for a representative image obtained from Untreated cell using the motion type classification 4-plots set. (D) Global view of all identified trajectories for a representative image obtained from Nocodazole treated cells, using the motion type classification 4-Plots set. (E) Comparison of the SMSS values frequency (i.e. relative frequencies expressed as percentages) distribution obtained with Untreated vs. Treated cells.

Despite significant advances (Martin et al., 2002; Gloter and Hoffmann, 2007) the effect of positional uncertainty and sample size on motion type estimates remain difficult to theoretically predict. Thus, we reasoned that a better approach would be to empirically estimate the uncertainty associated with each ODC and SMSS measurement (i.e., local error analysis). For this purpose, we developed a numerical method, based on the Monte Carlo simulation of artificial trajectories (Supplemental Information 1; Rigano and Strambio-De-Castillia, 2018b; Helmuth et al., 2007), whose true position with respect to the imaging system, rate of displacement (i.e., ODC), freedom of movement (i.e., SMSS) and length and are fully known (Rigano et al., 2018). After simulating the effect of positional error on these “ground truth” trajectories under different image quality contexts (i.e., SNR), ODC and SMSS are back-computed from the resulting “noisy” trajectories and the comparison between input and output values is used to estimate the uncertainty associated with motion type estimation as a function of expected motion characteristics (i.e., ODC and SMSS), motion duration (i.e., trajectory length L) and image quality (i.e., SNR). Using this information, the empirically generated four dimensional matrices relating L, SNR, ODC and SMSS values with expected values of ODC and SMSS, are interrogated by linear interpolation to obtain the uncertainty value associated with each trajectory under study. In turn, these ODC and SMSS uncertainty values are reported both in tabular format and as confidence intervals on two-dimensional ODC vs. SMSS scatter plots, which forms the basis for motion type classification in OMEGA.

### 2.7 Data Browser: tracking data and metadata exploration, provenance, persistence, and dissemination

The Data Browser is the main data and metadata management gateway for OMEGA (Figure 2-12; Supplemental Information 1), and it provides an intuitive interface that allows users to navigate across the entire data and metadata chain from images to trajectories, segments, tracking measures, and motion analysis uncertainties. In addition to allowing the import of pre-computed analysis results as described in Section 2.1.2, the Data Browser facilitates the execution of the following: 1) interactive navigation and display of analysis output already present in OMEGA; 2) export selected analysis results to the FS for downstream analysis either during a future OMEGA session or using third party applications. In addition, OMEGA results can be saved to the dedicated OMEGA Analysis Results repository (Figure 3 and Supplemental Figure 2).

#### 2.7.1 Data navigation

At each step of the analysis workflow (Figures 1 and 4), the user can decide to compare results obtained using different parameter settings. In order to do so, the Data Browser plugin-Superclass, provides a hierarchical navigation structure in which each node contains sets of intermediate data (i.e., detected particles, trajectories, edited trajectories, and segments) and function as a branching point for a dependent tree of possible downstream analysis results. Consistent with this tree structure, plugins that extend the Data Browser super-class, such as the currently available OMEGA Data Browser, facilitate the interactive display of dynamic lists that are populated with the analysis children of any selected data element (Figure 2-12; Supplemental Information 1). These resulting data trees are presented to the user using a familiar column view interface, where relevant metadata and result summaries are displayed at the bottom of each column to facilitate the identification of the desired data path. Additionally, to reduce work space clutter, results that at any point of time are not of immediate interest can be temporarily hidden from the view by unchecking their selection mark. Once identified and selected, results branching off a given element can be displayed in parallel on all concurrently opened results visualization windows (e.g. Side Bar, Trajectory Browser, and Tracking Measures), or stored as described below.

#### 2.7.2 Automated tracking of data provenance

Information describing the “origin” and “lineage” of data is essential for scientists to be able to correctly interpret image analysis results. Such information is often referred to as describing “data provenance” and can be conceptualized as metadata capable of answering key questions describing manipulation events occurring during the data lifecycle (Ram and Liu, 2009). To facilitate the tracking of data provenance, the comparison of results across laboratories, the repeatability of analysis processes, and the reproducibility of results, OMEGA ensures the persistence of analysis results and of data provenance metadata by creating links that bridges between dedicated image data and metadata stores (Goldberg et al., 2005), and results management repositories (i.e., the FS or results databases), to store and subsequently mine analytical output (i.e. including source image metadata, particle positions, trajectories, motion analysis results, associated uncertainties and analysis definition parameters). In either case, data is structured using a schema that recapitulates MIAPTE guidelines (Rigano and Strambio-De-Castillia, 2017; 2016). The use of MIAPTE facilitates management of data quality, particle tracking, motion analysis and error estimation results and facilitates meaningful comparison and reproduction of results obtained at different moments in time and from different laboratories.

#### 2.7.3 Persistence of tracking and motion analysis results

To ensure data and metadata persistence and downstream processing in future OMEGA sessions or third-party applications, particle tracking and analysis results data chains and related metadata describing their provenance can be saved to the dedicated OMEGA Analysis Results repository or exported to the FS using the OMEGA Data Browser export functionality (Figures 1 and 3 and Supplemental Figure 2). Using this functionality the user can choose to export specific results or entire OMEGA particle tracking and trajectory analysis sessions in order to resume work at a later time. Additionally, downstream statistical analysis or meta-analysis can be performed using third party applications (i.e., R or Matlab). Finally, data can be exchanged with other researchers for results comparison and reproduction.

## 3 Example use-cases and applications

We present here two use cases to illustrate OMEGA functionality. The first test case takes advantage of simulated image datasets that were produced to directly compare different SPT algorithms (Chenouard et al., 2014); scenario IV, infecting viral particles; SNR = 7; low particle density). As expected, when images were subjected to SPT within OMEGA, most trajectories displayed a stretched out appearance with occasional direction transitions, mimicking a condition where most particles display the tendency of “flying” over long distances in a particular direction (i.e., Lévy flights; Levy, 1937), such as what is observed in active motion (Figure 7). When trajectories were manually inspected, most appeared to be correct. However, trajectory nr. 424 appeared to be composed of two erroneously linked trajectories (Figure 7B), consistent with the observed “kinked” shape of the MSS curve (Figure 7C, bottom left, *arrowhead*). The use of the Trajectory Editing tool (Figure 7 A and D) allowed to split this trajectory in two individual trajectories, 424.1 and 424.2 (Figure 7E and F). While trajectory 424.1 gave rise to a straight MSS curve for trajectory consistent with the identification of a uniformly mobile particle, a similar analysis of 424.2 produced a bent curve indicating that such trajectory is apparently produced by a particle whose mobility is non self-similar and would require segmentation (Figure 7F). After trajectory editing, all resulting trajectories were subjected to diffusivity analysis (Figure 7G-K). The results of such analysis were consistent with the trajectories displaying directed motion, as indicated by the clustering of trajectories in the top quadrant of the phase space (Figure 7J, *red circled area*) as well as the prevalence of SMSS values close to 1 (Figure 7K). In particular trajectories nr. 27 and 32, as well as edited trajectory nr. 424.1, displayed a straight MSS curve consistent with uniform mobility. It should be noted that the ODC and SMSS estimation errors we observed are consistent with a high SNR level and relatively short trajectories (i.e., relatively small ODC error and relatively high SMSS error).

The second OMEGA test case is a real-life example provided by the Hunter’s lab (Clark et al., 2013; Pereira et al., 2012). In this example, CMMT rhesus macaques (*Macaca mulatta)* mammary tumor cells chronically infected with Mason-Pfizer Monkey Virus (M-PMV), a D-type retrovirus, were co-transfected with a plasmid expressing a codon-optimized GFP-tagged variant of the M-PMV Gag precursor polyprotein alongside one expressing mCherry-Tubulin (i.e., a microtubule subunit). Seven hours post-transfection cells were either mock-treated (Figure 8, *Untreated*) or treated with the microtubule polymerization inhibitor Nocodazole (Figure 8, *Treated*) for 1 hour prior to live imaging to observe the assembly of viral particles and their trafficking towards the plasma membrane. Example images were first imported into OMERO and then loaded into OMEGA for SPT (Figure 8A and B). Trajectories were examined using the OMEGA DTM plugin and a subset of trajectories displaying uniform mobility as indicated by a straight MSS graph, were assigned a specific motion type as indicated by the position of the line on the MSS plot. In Untreated cells, most trajectories were found to display a sub-diffusive behavior (Figure 8B and C, *fuchsia*) with a minority of viral particles displaying clearly diffusive and super-diffusive mobility (Figure 8B and C, *blue* and *purple* respectively). An example super-diffusive trajectory (i.e., *purple*) is displayed in the bottom insert in panel 7B. As expected, when cells were treated with Nocodazole, the global intracellular mobility of viral particles was dramatically reduced as testified both by the overall collapse of resulting trajectories (Figure C and D, compare top left panels) and by the clustering of viral trajectories in ODC vs. SMSS phase space (Figure 8C and D, right panels, compare the grey area vs. the red area). While the biggest effect was observed on ODC values, which drastically diminished, a significant effect was also observed on the MSS behavior as shown by comparing the resulting SMSS distributions in Untreated vs. Treated cells (Figure 8C and D, bottom left panels; Figure 8E). Of note, this analysis took a total of 5 minutes and required no manual tracking testifying the advantage of using OMEGA for increasing the throughput of systematic motion analysis experiments to higher levels than allowed by the use of individual and analysis tools.

## 4 Discussion

Despite tremendous improvements in space and time resolution of modern microscopic techniques, the translation of such advances towards increased understanding of intracellular particle movement has proceeded at a significantly slower pace. In order to circumvent this obstacle, one key aspect will be the development of shared infrastructure to foster collaboration between experimental scientists that have a deep understanding of the biological domain, and image analysis experts, mathematicians, statisticians, algorithm developers and software engineers that can help organize and make sense of the data. Ad hoc collaborations of this kind are increasingly becoming the norm between individual well-funded laboratories and occasionally at the inter-institutional level. However, in order to, facilitate the integration of genomics, transcriptomics, proteomics and functional data, and foster a systematic understanding of intracellular trafficking, it is necessary to build virtual “tables” where all required expertise can convene across time, space, and experimental systems. In order to provide a significant contribution towards this goal we have developed OMEGA, which was designed with the explicit goal to enhance reproducibility of particle tracking experiments.

Open-source, bioimage informatics initiatives largely focus on the production of general tools to address a great variety of analytical needs. In this context because of their flexible and extendable design, their user interface is generic and sometimes overwhelming.

In addition, while some of these available image processing systems provide integration with image data repositories as data sources and sinks (Taubert and Bucker, 2017; Carpenter, 2017; OME Consortium, 2018a; b; KNIME consortium, 2018; Walter et al., 2017), these tools do not bridge between processing and storage, do not offer support for automatic data provenance harvesting, and do not track error propagation (Table I; Tinevez et al., 2016; de Chaumont et al., 2012; Cardona and Tomancak, 2012).

We reasoned that a better approach would be to tackle a very limited set of biological questions and address them holistically from data acquisition to results interpretation (Taubert and Bucker, 2017). As a test case, we decided to study the dynamic behavior of retroviral viral particles during the initial phases of the viral life cycle for the following reasons:

1. <u>Realm of expertise</u>: The choice of this domain of study was motivated principally by the fact that retrovirus cell biology is well within our realm of expertise (Xu et al., 2013; Pertel et al., 2011b; a; Sokolskaja et al., 2010; Neagu et al., 2009; Sebastian et al., 2009; Strambio-De-Castillia and Hunter, 1992)
2. <u>Clear need</u>: Retroviral particle tracking is a relatively technology-poor domain of study, with well-documented and urgent scientific and quantitative analysis needs, and consequent opportunity for development.
3. <u>Opportunity</u>: SPT and motion analysis entail a well-defined series of analytical steps, which currently are not well integrated among one another and lack, metadata and procedure standardization as well as error estimation. Possibility of further collaboration: initiatives to foster community efforts for the improvement of SPT and motion analysis tasks are well underway (Chenouard et al., 2014).

The following design features of OMEGA set it apart from other platforms:

1. <u>Narrow focus and ease of use</u>: Because of its clear focus on the domain of intracellular particle tracking and motion analysis OMEGA executes a relatively narrow scope of computational tasks with the consequence that extreme interface flexibility can be substituted by a rich and intuitive Java graphical user interface (GUI) that massively reduces the user learning curve and facilitates the execution of repetitive manual work for many standard steps thus freeing the user to pay more attention to the actual experiments.
2. <u>Modularity and interoperability</u>: OMEGA facilitates interoperability and extension by way of a plugin architecture with well-defined application programming interfaces in which the calculation logic responsible for each step of the workflow is separated from the execution logic, representing the overall process that executes all computational steps.
3. <u>Bridging between data processing and management</u>: In order to establish a persistent bridge between storage and processing, OMEGA automatically harvests data provenance metadata for each executed particle tracking workflow, and uses them to annotate calculation results. Such metadata comprises Universally Unique Identifier (UUID) assigned to each data element and each step of the analysis execution run, direct links with source images, plugin version control, and analysis definition parameter settings. OMEGA provides a rich navigation GUI called Data Browser to facilitate retrieval, perusal, and saving of full particle tracking data chains, including trajectory data, motion analysis results, error propagation and their complete provenance details.
4. <u>Minimum Information Standards</u>: In order to facilitate reproducibility the metadata definition data model adopted in OMEGA complies and extends existing standards such as OME-TIFF (Goldberg et al., 2005) and our recently proposed MIAPTE guidelines for particle tracking experiment reproducibility.
5. <u>Error propagation</u>: In order to provide a solid basis for the standardization of data sharing, OMEGA provides integrated and open-access support for uncertainty estimation and for the evaluation of how such error propagates through the analysis routine. This is arguably the first essential step towards true data standardization and data sharing.

## 5 Conclusions

OMEGA is a novel cross-platform data management system for particle tracking experiments. OMEGA is freely available, flexible and easily extensible. It links upstream image data and metadata with downstream motion analysis tools, it automates data handling, processing, quality monitoring and interpretation, ultimately facilitating the comparison of image analysis results, of data analysis routines and of uncertainty quantification both inside and across laboratories. OMEGA’s intuitive interface facilitates data selection, data import, analysis and reporting of analysis results and uncertainties. Data can be exported for use in third party tools and across laboratories laying the foundation for the meta-analysis of data generated by multiple users and making it possible, for example, to compare the effect of specific treatments on particle motion across different experimental systems.

In conclusion, OMEGA facilitates the cooperation of all players whose role is required to understand complex biological systems: biological scientists, image analysis experts, algorithm developers, statisticians and software engineers. OMEGA is developed following a modern open development paradigm, which allows the entire bioimage informatics community to participate in its development. Thus, OMEGA facilitates the process of incorporating both novel and already available tools to build integrated data processing and analysis pipelines for quantitative, real-time, sub-cellular particle tracking. By directly addressing issues of error propagation and data provenance, and by relying on semantic data models to record tracking results and analysis-definition metadata (Goldberg et al., 2005; Rigano and Strambio-De-Castillia, 2017; 2016), OMEGA lays the ground for the development of analytical standards for particle tracking. Such standards are a prerequisite for data reproducibility, reusability and transferability between research groups. Because of its ease of use, interoperability and extensibility, OMEGA could function as a proof-of-principle for the development of analytical, data management and collaborative standards that can be extended across diverse scientific questions, model systems and experimental contexts.

## Supporting information

07_Supplemental Table III

## Abbreviations

DGT: distance from ground truth
MIAPTE: minimum information about particle tracking experiments
SPT: single particle tracking
MPT: multiple particle tracking
MSD: mean squared displacement
MSS: moment scaling spectrum
ODC: observed diffusion constant
OMERO: open microscopy environment remote objects
SNR: signal to noise ratio
TS: Trajectory Segmentation

## 6 Acknowledgements

The success of this multidisciplinary project was due to the collective effort of many individuals. We want to specifically acknowledge our debt of gratitude toward those without whom this software would simply not have been produced. We are deeply beholden to **Peter Kunszt** (SystemsX.ch now at Dynatrace Barcelona) for continual encouragement, critical support (financial and otherwise), and lots of helpful discussions. <u>Without Peter this project would never have started!</u> We thank **Andrea Danani**, **Roberto Mastropietro** (University of Applied Sciences and Arts of Southern Switzerland - SUPSI) and **Bernd Rinn** (ETH – Zurich), for their assistance and guidance, for invaluable discussions, and for their generosity with their software development and project management expertise during pivotal times along our path. We thank **Orlando Petrini** and **Mauro Tonolla** (Istituto Cantonale di Microbiologia, Bellinzona, Switzerland) for active hospitality, helpful discussions and continual morale-boosting to C.-S.-D.-C. We are extremely grateful to **Karl Bellvé**, **Kevin Fogarty** and **Lawrence Lifshitz** (University of Massachusetts Medical School) for their support and encouragement, their generosity with code, time, knowledge and experience, and for critical discussions and feedback. We are indebted to **Jay Copeland**, **Mario Niepel** and **Peter Sorger** (Harvard Medical School), **Kevin Eliceiri** (LOCI, University of Wisconsin at Madison), **David Grunwald** (University of Massachusetts Medical School, Worcester, MA), **Martin Spitaler** (Max Planck Institute of Biochemistry - Munich), **Jason Swedlow** (University of Dundee) and **Mark Woodbridge** (Imperial College London) for stimulating discussions, inspiration and motivation. We are immensely grateful to **Diego Frei**, **Loris Grossi** (University of Applied Sciences and Arts of Southern Switzerland - SUPSI), **Pietro Incardona**, **Krysztof Gonciarz** (Max-Plank Institute of Molecular Cell Biology and Genetics - Dresden), Lawrence Lifshitz and **Curtis Rueden** (LOCI, University of Wisconsin at Madison), for sharing their code and software engineering skills with A.R. We thank the entire **Open Microscopy Community** (openmicroscopy.org) for inspiration, encouragement, motivation and help. They have been our role models throughout and as such their contribution is simply too extensive to be properly documented. Our gratitude extends but is not limited to: **Chris Allan**, **Sebastian Besson**, **Jean-Marie Burel**, **Melissa Linker**t, **Josh Moore**, **Will Moore**, and Jason Swedlow (OME and Glencoe Software), **Christian Dietz**, Kevin Eliceiri and Curtis Rueden (ImageJ/Fiji and KNIME ecosystem). We are grateful to Kevin Fogarty and Kevin Eliceiri for critically reviewing the manuscript. We apologize to colleagues whose work we were not able to cite due to space limitations.

Extramural funding was from the Swiss National Science Foundation (Project CRSII3_136282 to C.-S.-D.-C. and J.L.), the European Commission FP7 (Project HEALTH-2007-2.3.2, GA HEALTH-F3-2008-201,032, to C.-S.-D.-C. and J.L.) and the National Institutes of Health (National Institute on Drug Abuse Project 5DP1DA034990 to J.L..) Institutional funding was from the Institute of Research in Biomedicine and the University of Geneva (to C.-S.-D.-C. and J.L.), **SystemsX.ch Information Technology**, **Emory University** (to A.R., R.G. and V.G.), **University of Applied Sciences and Arts of Southern Switzerland - SUPSI** (to A.R., R.G. and V.G.), and the **University of Massachusetts Medical** school (to A.R., C.-S.-D.-C., J.L. and V.G).

## Supplemental material

**<u>Supplemental Information 1</u>** – Benchmarking, validation and software architecture

**<u>Supplemental Information 2</u>** - getMSS_pseudocode.txt

**<u>Supplemental Information 3</u>** – calcDiffusion_pseudocode.txt

## Supplemental Figures

**<u>Supplemental Figure 1:</u> Typical OMEGA dataflows**

**<u>Supplemental Figure 2:</u> OMEGA for developers: logical structure and software architecture s**

**<u>Supplemental Figure 3:</u> Uniform artificial trajectories examples**

**<u>Supplemental Figure 4:</u> Validation of the SNR Estimation plugin**

**<u>Supplemental Figure 5:</u> Comparison of getMSS with alternative ODC estimation methods**

**<u>Supplemental Figure 6:</u> Validation of getMSS over extended Brownian trajectories test sets**

**<u>Supplemental Figure 7:</u> Validation of OMEGA motion type estimation**

**<u>Supplemental Table I:</u>** Mathematical Analysis of Motion in OMEGA

**<u>Supplemental Table II:</u>** Motion type classification criteria

**<u>Supplemental Table III:</u>** Benchmarking test cases

