## Supplementary material for "OMEGA: a software tool for the management, analysis, and dissemination of intracellular trafficking data that incorporates motion type classification and quality control": 01_Supplemental Information 1

#### Table of content

**Supplemental Information 1** - this document

**Supplemental Information 2** - getMSS_pseudocode.txt

**Supplemental Information 3** - calcDiffusion_pseudocode.txt

**Supplemental Figures**

**Supplemental Figure 1: Typical OMEGA dataflows**
**Supplemental Figure 2: OMEGA for developers: logical structure and software architecture s**
**Supplemental Figure 3: Uniform artificial trajectories examples**
**Supplemental Figure 4: Validation of the SNR Estimation plugin**
**Supplemental Figure 5: Comparison of getMSS with alternative ODC estimation methods**
**Supplemental Figure 6: Validation of getMSS over extended Brownian trajectories test sets**
**Supplemental Figure 7: Validation of OMEGA motion type estimation**

**Supplemental Table I:** Mathematical Analysis of Motion in OMEGA

**Supplemental Table II:** Motion type classification criteria

**Supplemental Table III:** Benchmarking test cases

### Supplemental Information 1

#### *System overview and computer implementation*

OMEGA (Figure 1; Rigano and Strambio-De-Castillia, 2018a) is written entirely in Java (viz. Java 8) and uses external libraries including: *SWT* for the GUI, *OMERO.Blitz* (including *Ice* and *Hibernate*) and *MySQL Connector/J* for communication with databases, *imageJ* (including *ij2util,* , *imglib2*, *SciJava* and *SCIFIO*) for image processing, the *MOSAICsuite* for tracking (Sbalzarini and Koumoutsakos, 2005; Incardona and Sbalzarini, 2014), and *JFreeChart* for plotting (OME Consortium, 2017a; SWT consortium, 2018; OME Consortium, 2017b; Object Refinery Inc, 2017; SQL project, 2018; MOSAIC group, 2006; SciJava consortium, 2017). To enhance usability by experimental biologists regardless of their level of computational proficiency, the OMEGA information system relies on graphically supported user interactions mediated by a rich human-machine graphical user interface (GUI; Figure 2). This GUI mediates all interactions with the application and facilitates the execution of all aspects of the analysis workflows in the context of information delineating the provenance of data. OMEGA interacts with the OMERO image data repository (Allan et al., 2012) to retrieve image data and metadata to be subjected to analysis and with a dedicated analysis results repository to store and mine analytical output (Figure 3, and Supplemental Figures 1 and 2). Particle tracking is highly dependent on image quality, noise level, and experimental conditions (Chenouard et al., 2014; Jaqaman and Danuser, 2009), making it often necessary to test different algorithms before deciding which works best for a specific condition. To facilitate this task, OMEGA is designed to allow stringing independent tracking algorithms together in a modular fashion, while at the same time offering generalized mechanisms for handling parameterization, analysis results and data provenance (Figure 3 and Supplemental Figure 2). Thus while the conceptual data path flows from one analytical module to the next (Figures 1 and 3 and Supplemental Figure 1), in reality each modular element receives input independently from and sends output directly to, the functional core of OMEGA (Figure 3 and Supplemental Figure 2, *Core*). This architecture facilitate the extension of the analysis toolbox as it provides a mechanism for linking together plugins regardless of the specificity of their inner mechanisms, provided the output of one element matches the input of the next. To make this possible, OMEGA takes advantage on semantic data typing as sanctioned by the OME-XML (Goldberg et al., 2005) and by our recently proposed MIAPTE data models to define the structure, meaning and behavior of data and metadata stored and manipulated by the system (Rigano and Strambio-De-Castillia, 2016a; b; 2017b)..

To facilitate the production, troubleshooting, interpretation, dissemination, re-use and reproduction of experimental viral tracking and trajectory analysis results across multiple laboratories, and in order to promote the interaction between bench scientists, theoreticians, mathematicians and software engineers, OMEGA complies with guidelines that have been proposed for the development of open-source scientific software (Cardona and Tomancak, 2012; Carpenter et al., 2012). Thus, in our design decisions emphasis was given to usability for both users and developers, to data provenance recording , to full integration of work- and dataflows, to interoperability, to the use of validated and well documented analysis routines, and last but not least to the estimation and tracking of measurements uncertainty.

#### OMEGA for users: Graphical User Interface-based interaction

The GUI of OMEGA takes advantage of components that are commonly used by other applications, such as: 1) the main workspace window for managing workflows, interacting with plugins, and inspecting results. 2) The top toolbar (Figure 2-1) containing individual 'button' elements to launch the plugin-selection window each analysis modules is linked with. 3) Plugin launcher windows (Figure 2-2) used to select which plugin the user intents to launch. In addition to these familiar elements, OMEGA contains six distinctive graphical components, which are detailed below.

##### OMERO image browser

The OMERO Image Browser (Figure 2-3) currently available in OMEGA utilizes a minimal navigation interface that allows the user to obtain relevant image metadata while at the same time avoiding overcrowding and limiting undue duplication of functionalities already available elsewhere. This window takes advantage of the OMERO Project, Dataset, Image hierarchical structure (Goldberg et al., 2005; Allan et al., 2012) to navigate across data and display all content belonging to a specific user in either a list or grid view. Only image thumbnails and metadata essential for image selection are shown while additional viewing options are delegated to the OMERO (Allan et al., 2012).

##### Sidebar

A minimal image viewer positioned on the Side Bar is responsible for displaying image data and metadata after they have been loaded from an available OMERO database (Figure 2-4, 6 and 10). To facilitate browsing among loaded image data, the side bar also shows annotations related to datasets and projects used to organize image data in the source image repository. Last but not least, the sidebar has also a key role in trajectory inspection. Upon completion of particle detection and linking, trajectory editing and segmentation, resulting particles, trajectories and segments can be selected using drop down menus and overlaid over the corresponding image-plane displayed on the sidebar to produce a visual impression of the extent of motion of each individual particle. Trajectories can be displayed either in their entirety or dynamically depending on the displayed time frame.

##### Plugin window

The general structure of the pane is shared by all plugins that utilize this graphical pane for user interactions and consists of: 1) a data navigation area; 2) an execution queue area; and 3) a panel for analysis definition and results inspection (Figure 2-5). Thus, when OMEGA contains data that is appropriate for a specific plugin (e.g. images to be subjected to particle detection) such data appears in the "Loaded Data" area of the window, ready for selection by the user. Once the analysis run has been defined, the user can add it to the execution queue by pressing the "+" button. Once all desired runs have been loaded, their execution can be launched in the background (i.e. allowing the user to continue working on other tasks) and monitored using the status bar located at the bottom of the window. While the run definition and result inspection panels are designed to respond to the needs of each individual plugin, they share a common general organization to expedite user familiarization and minimize the learning curve. Thus the "Run Definition" tab is organized in four quadrants, which proceeding left to right and top to bottom contain information describing the input data (i.e. "Input Information"), data selection parameters (i.e. "Selection Parameters") and algorithm specific parameters including advanced options when required.

At the end of each run, output results become available for inspection by selecting the desired execution run and selecting the "Results" tab. The results tab consists of a generic results summary area, which reports general information such as user name, execution date and aggregated results, as well as of an algorithm specific results area.

Particle detection

The image selection tree (Figure 2-5) facilitates the selection of images to be analyzed. Once a selection has been made, the "Run Definition" tab allows the user to visualize image details, enter physical dimensions (i.e. pixel and voxel sizes and time per frame), when none have been automatically imported from OMERO, select a specific z-plane and channel to be subjected to analysis and set algorithm specific parameters. At time of writing the only available plugin for particle detection is the *MOSAICsuite* Feature Point Detection (FPD) plugin, whose *Radius*, *Cutoff* and *Percentile* parameters can be set following published specifications (Sbalzarini and Koumoutsakos, 2005; Incardona and Sbalzarini, 2014). The "Detection Results" tab displays general run results as well as results associated with each particle as defined by the algorithm developer and by MIAPTE requirements (Rigano and Strambio-De-Castillia, 2016a; b; 2017b).

Particle linking

The data selection tree of this plugin allows the selection of the particle detection run to be. Once a specific particle detection run has been selected, the "Input Information" quadrant of the "Run Definition" tab displays relevant details of that run in order to facilitate selection. After selection, the user can set both basic and advance parameters when available. At time of writing the only particle linking plugin available in OMEGA is the *MOSAICsuite* Feature Point Linking (FPL) (Sbalzarini and Koumoutsakos, 2005; Incardona and Sbalzarini, 2014), which requires the *Link Range*, *Displacement*, *Movement Type*, *Object Feature*, *Dynamics*, *Optimizer* and *Minimum Trajectory Length* parameters to be set by the user. In addition to presenting general run results, the "Linking Results" panel also presents tracking results where the position of each particle in a trajectory is associated with trajectory name and with other MIAPTE-compliant information (Rigano and Strambio-De-Castillia, 2016a; b; 2017b).

SNR estimation

Using this plugin the user can use the top "Select image" data selector to select the desired image and the "Loaded Data" selector to choose an associated Particle Detection run to be analyzed. Subsequently, the user can set the appropriate parameters for the run. Information regarding both images and detected particles is available in the "Input Information" area of this window to facilitate selection. The results section is subdivided in three tabs depending on the granularity of the required results. The "ROI results" tab reports results associated with each particle while the "Plane results" and the "Image results" tabs present measures aggregated by plane and by image respectively.

**Trajectory Browser**

This graphical element is available in all Trajectory Manager and Trajectory Measures associated plugins (Figure 2-9), and facilitates the user interactions needed to edit, segment and analyze the motion characteristics of each trajectory or trajectory segment. The pane is associated with a data selector top bar, which allows the user to select the desired image and associated detection, linking, editing, segmentation and trajectory measures runs (as appropriate) using a series of nesting combo boxes. Once a specific image / run combination has been selected, the browser paints all associated trajectories in the form of a sequence of linked blocks, which depending on user needs might contain individual particle thumbnails. Finally, individual trajectories can be selected among all available for visualization as image overlays and for further manipulation.

Trajectory editing

This plugin facilitates the manual inspection and curation of particle linking results that originate from automated tracking. The trajectory editing plugin utilizes the Trajectory Browser graphical pane twofold. After selecting a linking run, the "Browser" tab aides the selection of one or more trajectories to be subjected to re-linking. Once individual trajectories have been selected, the "Editor" tab employs a modified version of the Trajectory Browser GUI to split and merge trajectories utilizing facilities made available via the right-click menu.

Trajectory segmentation

Trajectories of diffraction-limited objects moving within a cell are often characterized by rapid shifts in direction and rate of motion as the object interacts with the cellular milieu. If a change of behavior is detected either by eye or thanks to the workflow depicted in Figures 5 and 6, the trajectory in question can be subdivided in segments of putative homogeneous mobility by utilizing the Trajectory Segmentation plugin. This plugin utilizes the Trajectory Browser for trajectory selection and a specialized Trajectory Segmentation panel (Figure 3-8) for subdividing the trajectory in segments and for annotating individual segments with one of five motion type labels (Supplemental Table II). This segmentation panel employs a dependent graphical component to display each selected trajectory on an x-y Cartesian space that mimics the associated image-plane and to select and annotate specific portions of it. Annotation can be executed in one of two modalities. In the "Select Spot" mode, the selection of the Start and End position of each segment is followed by Motion Type assignment. In the "Select Motion Type" mode, the selection of the desired Motion Type is followed by the selection of the segment extremities.

##### Tracking Measures

All available Tracking Measures plugins (Figure 4, Supplemental Table I and main text), utilize the same basic graphical format to facilitate user interactions (Figure 2-7.1, 7.2 and 11). In all cases, the window pane is subdivided in the following tabs: 1) the "Browser" is used to select one or more trajectory-segments belonging to a specific image / run combination; 2) the "Run" tab is used to execute specific tracking measures runs; and 3) the "Plots" tab is used to visualize analysis results in graphical form; and 4) one or more "Results" tabs present numerical results in tabular format following MIAPTE guidelines (Rigano and Strambio-De-Castillia, 2016a; b; 2017b). The "Plots" tab is organized in a similar fashion for all Tracking Measures plugins. Namely, it consist of a left side area where a series of combo-box selector elements allow to choose between local and global measures, type of measure, and type of graph. Once the selection has been made the desired plot is displayed on the right side of the pane. Three types of graphs are allowed, line, bar and histogram distribution.

##### Data browser

The Data browser (Figure 2-12) dynamically displays all analysis output present in OMEGA in a column view that facilitates data tree navigation and selection. Specifically designed check-boxes can also be used to temporarily hide specific data branches from the user when necessary to avoid clutter. To facilitate the selection of desired results paths, relevant metadata and summary results pertaining to each selected element are displayed at the bottom of each column. Using this GUI element, pre-computed particles or trajectories can be imported in the application and associated to Orphaned analyses data elements to be subjected to further analysis. Finally, in the OMEGA Data browser particles, trajectories and analysis data produced by third party application can be imported, and OMEGA results selected and exported in various file formats to the local file system (FS).

#### OMEGA for developers: Logical structure and software architecture

OMEGA is composed of three structural elements that work in concert with five peripheral components to communicate with external data stores and with the FS (FS; Supplemental Figure 2). The three structural elements are: Omega/Core (Rigano and Strambio-De-Castillia, 2018a), OmegaCommons Application Programming Interface (API) and OmegaCommons/Analysis plugin-Superclasses (Rigano and Strambio-De-Castillia, 2018b). The five communication components comprise: 1) the OmegaOmeroCommons/Image Gateway that is required for OMEGA to interact with the Image Repository (Rigano and Strambio-De-Castillia, 2018c); 2) the Omega/Core Analysis results Gateway, which is responsible for communicating with the Analysis Results Repository; 3) the Log Manager; 4) the Configuration Manager; and 5) the Analysis Import/Export Manager .

##### Omega/Core

The Omega/Core (Rigano and Strambio-De-Castillia, 2018a) of the application contains all main sub-components including the application launcher, the initialization and disposition methods and event-driven logic driving all communication between the core and the analysis plugins. The Omega/Core also contains all GUI sub-components including the top toolbox, the side bar and the workspace, the analysis results repository preferences window, and the plugin launcher window, which is opened every time users click on a module button on the toolbox. Moreover, the Omega/Core contains facilities to handle both Log and Configuration managers.

#### OmegaCommons

The OmegaCommons API (Rigano and Strambio-De-Castillia, 2018b) was designed to facilitate the extraction of OMEGA libraries for custom plugin development. In addition to general utility methods, it includes Java classes used as building blocks for the construction of individual plugins, interfaces necessary to mediate the communication between plugins and the Omega/Core, and the system used for event-driven communication. Commons API also contains all foundational libraries necessary for trajectory analysis in OMEGA. Such libraries are composed of math-oriented static Java classes, are sub-divided in five categories based on the tracking measure they are responsible for (i.e. Intensity, Mobility, Velocity, Diffusivity and Diffusivity Measures Uncertainty Estimation), and are specifically designed to facilitate the generation of stand-alone Tracking Measures tools. Generic GUI elements, as well as common constants and exceptions were included here to facilitate the application accessibility by third-party plugins.

In addition, Commons API contains three categories of data structures that are designed to store image data and metadata as well as the results produced at run time by the OMEGA application. The first group includes structures designed to contain data and metadata originating from OME-XML specific elements, such as Project, Dataset, Image, Experimenter and Experimenter Group. The second group of data structures was created on the basis of the OMEGA data model and MIAPTE specifications (Rigano and Strambio-De-Castillia, 2016a; b; 2017b) to store data produced as output by the analysis pipeline. Since most of OMEGA data-producing processes require parameter input, these data structures are constructed to allow the storage of both results and parameter settings associated with each analysis runs. The third group contains generic data structures whose purpose is, for example, to store data necessary to establish connections with remote data stores, such as server information (i.e. hostname and port) and user credentials (i.e. username, encrypted password). This arrangement allows developers to extend or implement each components thereby facilitating the development of third-party plugins for connecting OMEGA to servers other than OMERO and OMEGA.

#### Analysis plugin-Superclasses

The main business logic in OMEGA is organized around six modular plugin-Superclasses: 1) Image browser; 2) Particle tracking; 3) SNR estimation; 4) Trajectory manager: 5) Tracking measures; and 6) Data browser (Figure 3 and Supplemental Figure 2, solid lines boxes). These types are extended by one or more interchangeable plugins, which in turn are responsible for specific steps of the analysis data-stream. Currently, OMEGA ships with a set of two Data Management (Figure 3, *orange boxes*), nine Analysis (Figure 3, *blue boxes*), and one Quality Control (Figure 3, *green boxes*) plugin. The Data Management plugins are: 1) OMERO Image browser; and 2) OMEGA Data browser. The Analysis plugins are: 1) *MOSAICsuite* FPD; 2)

*MOSAICsuite* FPL; 3) *MOSAICsuite* Feature Point Tracker (FPT); 4) OMEGA Trajectory Editing; 5) OMEGA Trajectory Segmentation; and 6-9) OMEGA Intensity-, Mobility-, Velocity- and Diffusivity- Tracking Measures. The only Quality Control plugin currently available is MOSAIC SNR Estimation.

It is expected that further plugins will be developed, either by the OMEGA team or by third parties. Following extension policies that are being developed, each plugin will be required to provide its own GUI panel, to extend its generic Java Class and to implement Java interfaces in order to exchange data from the application. Currently each plugin is placed within its own Java package and is an integral part of the application. However, OMEGA's design allows them to be extracted so that, at later development stages, they could be loaded independently and allow the application to handle both native and third party plugins.

Image Browser

The only currently available plugin that extends this super-class is the OMERO Image Browser. This plugin implements an ImageJ2 functional operation (ImageJ2 Ops) we recently developed for the OMERO-ImageJ initiative (Rigano and Strambio-De-Castillia, 2017a), to access image data and metadata stored in available OMERO servers. Specifically, this ImageJ2 Op takes advantage of the OMERO API to provide a gateway that mediates all interactions with the OMERO server and provides a middle layer of data structures where image metadata is transitorily stored. When the Experiments selects an image to analyze, this triggers events whereby image pixels and metadata is loaded into OMEGA. The process to load images from the OMERO database is implemented as a separated thread with respect to the GUI in order to increase its concurrent responsiveness. While facilities to import image data and metadata from OMERO ship with OMEGA, by providing interfaces that can be adapted to allow the communication with other image data sources, the software is not exclusively restricted to interacting with OMERO.

Particle tracking

The identification of feature points and trajectories in OMEGA is carried out by plugins associated with the Particle Tracking super-class (Figures 1, 2-5 and 3). This plugin type is compatible with three alternative plugin designs: 1) Particle Detection stand-alone; 2) Particle Linking stand-alone; and 3) integrated Particle Tracking plugin merging both detection and linking functionalities in a single element. Using this strategy, OMEGA can incorporate a large variety of tracking algorithms regardless of their specific implementation style. Furthermore, this arrangement allows the user to join particle detection and linking algorithms originating from different sources increasing the probability of finding a successful combination (Chenouard et al., 2014).

Once started, the Particle Detection's main-thread launches a dependent thread for each image-plane, collects results from each of these threads, and then transfers them back for display by the GUI. Once the process is finished the master thread is in charge of packaging the information received and of sending events to the application core for further processing. Particle Linking plugins work much of the same way except they consist of single thread processes.

At time of writing OMEGA incorporates the *MOSAICsuite* ImageJ plugin implementation of the Sbalzarini and Koumoutsakos algorithm (Sbalzarini and Koumoutsakos, 2005; Incardona and Sbalzarini, 2014). Because this plugin was originally implemented as a dual-function particle tracker, in addition to creating a plugin incorporating the original *MOSAICsuite* version (Incardona and Sbalzarini, 2014), the algorithm was subdivided into its particle detection and linking components. This gave rise to three independent *MOSAICsuite* FPD, *MOSAICsuite* FPL and *MOSAICsuite* FPT OMEGA plugins, which are part of the current OMEGA package.

Trajectory Manager

Plugins defined by this super-class, utilize the Trajectory Browser graphical element (Figure 2-9), to facilitate inspection and editing of individual trajectories. These plugins generate one main thread for both data display and manipulation and a separate thread that handles particle thumbnails loading. Currently this type is extended by two plugins: Trajectory Editing and Trajectory Segmentation.

Tracking Measures

The Tracking Measures super-class defines plugins that compute, retrieve, and display tracking measures and estimation error associated with each particle, link, trajectory or individual segment of interest. At the time of writing, they include: OMEGA Intensity, Mobility, Velocity, and Diffusivity Measures. An additional Anisotropy Measures plugin is under development. Whenever possible results handled by plugins extending this modular super-class are automatically computed by the application core on the basis of particles and trajectories obtained from the tracking plugins. Additionally, mathematical functions that require parameter settings are executed *ad hoc* from within each plugin. In all cases, tracking measures plugins use the Jfreechart Java library to plot trajectory analysis results for inspection, interpretation and presentation.

SNR Estimation

The only plugin associated with the SNR Estimation type currently available in OMEGA utilizes a custom Java implementation of the *MOSAICsuite*'s Local SNR Estimation algorithm originally developed by the MOSAIC group in Matlab (Rigano et al., 2018; Gong and Sbalzarini, 2016; Xiao et al., 2016). This plugin estimates the local SNR of each particle identified by the particle detector, computes global SNR values for all trajectories of interest, and calculates aggregate values at both the plane and the image levels. Plugins associated with this type generate a master thread upon input image and particle detection run selection, which in turn distributes work to multiple dependent threads that execute independently on each image-plane to be analysed.

Data Browser

This modular super-class defines the OMEGA data browser plugin, which utilizes the Data Browser graphical element (Figure 2-12) to traverse tree branches that originate from each image accessible by the application and connect them to each derived analysis leaf produced by the tracking pipeline. Consistent with the analysis workflow structure, plugins extending this super-class allow data browsing through dynamic lists that are populated interactively based on the user necessity to drill in on specific data elements.

##### Image Gateway

At the time of writing, the only available Image Gateway in OMEGA is the OmegaOmeroCommons/OmeroGateway (Supplemental Figure 2, *blue box*). This component implements and extends the *OMERO.blitz* library (Rigano and Strambio-De-Castillia, 2018c), and other components of the OMERO API to communicate with OMERO as the main source of images. Additional connectors will be developed as needed.

##### Analysis Results Gateway

The Analysis results Gateway (Supplemental Figure 2, *red box*) is a custom built middleware, which contains all the Java classes that are necessary for the Omega/Core to interact with both the OMEGA Analysis Results repository in order to read or write data packets represented by the OMEGA model. At the time of writing, in addition to utilizing standard Java libraries, it leverages the API offered by the *MySQL Connector/J* library (SQL project, 2018). Connectors to additional results repositories will be developed as needed.

##### Analysis Import/Export Manager

The Import/Export Manager (Supplemental Figure 2, *green box*) is responsible for handling all operations needed to read or write trajectory, motion analysis and uncertainty estimation data and provenance metadata from or to the FS respectively.

##### Log Manager

The Log Manager (Supplemental Figure 2, *green box*) records all exceptions across the application and saves them in a set of predefined log files using a policy that subdivides errors based on their origin. In addition, a general catch-all category is implemented for uncategorized error types.

##### Configuration Manager

The Configuration Manager (Supplemental Figure 2, *green box*) interacts with the configuration files that are written by the application on the local FS. The main task carried out by this manager is to retrieve and load options relative to each plugin at application launch, as well as to collect and write newly set option values when the application is closed.

#### Data Storage

The persistence of particle positions, trajectories, motion analysis results, analytical errors and associated metadata is made possible either by exporting data to the FS or by writing them to a dedicated OMEGA relational database implemented in MySQL. In both cases data is stored as specified by relevant portions of the OME-XML data model (Goldberg et al., 2005) as well as of the MIAPTE-compliant OMEGA data model (Rigano and Strambio-De-Castillia, 2016a; b; 2017b). All input and output operations are mediated by the OMEGA application making them transparent to the user.

#### *Validation and benchmarking*

Extensive validation of OMEGA components was conducted here and elsewhere (Rigano et al., 2018).

#### Validation of Particle Detection and Particle Tracking algorithms

Validation of the particle detection algorithm was performed as described (Rigano et al., 2018). The functionality of the integrated OMEGA plugins was compared with the stand-alone *MOSAICsuite* Java (i.e., ImageJ plugin) versions of the same code and results were found to be comparable (not shown).

#### Validation of SNR Estimation algorithm

The OMEGA SNR Estimation plugin uses a custom Java implementation of the *MOSAICsuite*'s Local SNR Estimation algorithm (Sbalzarini and Koumoutsakos, 2005; Gong and Sbalzarini, 2016; Xiao et al., 2016; Rigano et al., 2018) to compute the local SNR for each particle identified localized during particle detection, as well aggregate measures at the trajectory, plane and image level. This OMEGA Java implementation of the SNR algorithm was tested on a set of artificial images (Rigano and Strambio-De-Castillia, 2018d; Rigano et al., 2018) each containing moving point sources at different know local SNR and generated as previously described (Sbalzarini and Koumoutsakos, 2005; Rigano et al., 2018). After estimation of local SNR values according to Cheezum (Cheezum et al., 2001), the relative $SNR_{Cz}$ Distance from Ground Truth (DGT) was computed as follows:

$$Relative\ DGT_{SNR_{Cz}} = {(output\ SNR_{Cz} - input\ SNR_{Cz})}/{input\ SNR_{Cz}}$$

and corresponding absolute values were plotted as a function of input $SNR_{Cz}$ (Supplemental Figure 4). Observed $SNR_{Cz}$ values were in good agreement with published results (Sbalzarini and Koumoutsakos, 2005; Rigano et al., 2018), and were measured to within 10 % of ground truth for all $SNR_{Cz}$ values greater than 2.5 and to within 5 % for all values of $SNR_{Cz}$ greater than 3.0. In addition, as expected SNR estimation uncertainty was observed to converge towards zero with increasing SNR values. Overall, these results confirm that the functionality of the algorithm was not altered as a consequence of having been ported from Matlab to Java and incorporated into OMEGA.

#### Algorithms for the generation of artificial trajectories

##### Generation of artificial trajectories of known input ODC and SMSS

Artificial trajectories of arbitrary motion type and known ODC (viz. D in case of normal diffusive states) and SMSS were generated using the *TrajectoryGenerator* Matlab routine (Rigano et al., 2018; Rigano and Strambio-De-Castillia, 2018e), which is a modification of a previously published algorithm developed earlier for this purpose (Helmuth et al., 2007; Rigano and Strambio-De-Castillia, 2018e; Rigano et al., 2018). For validation purposes, we generated a test set composed of a total of 1936 test cases (Supplemental Table III).

##### Simulation of Brownian motion

In order to simulate Brownian motion of known input ODC (Supplemental Table III) to be used for benchmarking purposes (see below) we developed two algorithms specifically for this study (Rigano and Strambio-De-Castillia, 2018e). The first algorithm is called *BrownianTrajectoryGenerator* and was implemented in Matlab; the second, algorithm, termed *BrownianTrajectoryGenerator_2*, was implemented in

Perl (Supplemental Figure 5 *BrownianTrajectoryGenerator* and *BrownianTrajectoryGenerator_2*). Both *BrownianTrajectoryGenerator* and *2* were developed to generate random-walk trajectories of known length (L), ODC (input_ODC), and step interval (input_Δt). While, the two methods share the initialization steps, they differ in the method employed to generate trajectory points. Thus both *BrownianTrajectoryGenerator* and *2* algorithms begin with setting the first position of each trajectory to $P_0 = (0, 0)$ and by initializing two arrays to store the $x_i$ and $y_i$ coordinates of each subsequent point, $Pi = (x_i, y_i)$, of each trajectory. Subsequently, the expected Mean Square Displacement (MSD) based on input ODC and Δt is calculated (MSD = 2 * 2 * input_ODC * input_Δt). From this point forwards, the two algorithms differ:

1) <u>*BrownianTrajectoryGenerator*</u> calculates the positional vector of each subsequent point $Pi = (x_i, y_i)$, by randomly producing a random step size based on MSD and a random angle alpha and then combining them as follows:
    - xi = x(i-1) + step*cos(alpha)
    - yi = y(i-1) + step*sin(alpha);
2) <u>*BrownianTrajectoryGenerator_2*</u> calculates $Pi = (x_i, y_i)$ positions as follows:
    - xi → {if (random[0;1] < 0.5), xi = x(i-1) + MSD; else x = x(i-1) - MSD}
    - yi → {if (random[0;1] < 0.5), yi = y(i-1) + MSD; else y = y(i-1) - MSD}.

The only difference between the two methods is represented by the way in which the direction of motion is set. While in *BrownianTrajectoryGenerator* direction is determined by random generation of the angle and of the step size; in *BrownianTrajectoryGenerator_2* direction is determined by randomizing the sign of the x and y coordinates of the subsequent point.

#### <u>Algorithms for the estimation of ODC and SMSS</u>

In order to validate the ODC and SMSS estimation functions contained in OMEGA (see below) we utilized *getMSS* (Supplemental Figures 4, 5 and 6, *getMSS;* Rigano and Strambio-De-Castillia, 2018e), a Matlab function developed earlier for the same purpose (Sbalzarini and Koumoutsakos, 2005; Racine, 2011) and the two ODC estimation methods (Supplemental Figure 5, *log-log method* and *linear method*), present in *calcDiffusion* (Rigano and Strambio-De-Castillia, 2018e), a Perl script that was developed for this study (not shown). Both algorithms are summarized below. Pseudocode for *getMSS* and for *calcDiffusion* is provided in Supplementary Information 2 and 3 respectively.

##### getMSS

- Declare a two-dimensional <u>points</u> matrix, with index iP, that for each entry contains the paired (x, y) coordinates of points from input trajectories (e.g. from *BrownianTrajectoryGenerator*, *BrownianTrajectoryGenerator_2* or *TrajectoryGenerator*).
- Define <u>nMoments</u> as the number of moments of displacement of different order we wish to examine.
- Define <u>L</u> (length) as the number of points contained in the trajectory.

- Define L3 as the size of the Δt time interval (i.e. window) that will be used for subsequent computations. L3 is defined as L divided by a parameter that defines the divisor (e.g. the values of L3 employed in Supplemental Figures 5 and 6, were L/3, L/5, L/10 and L/250).
- Define an array called frameshifts (with index iFShift) containing numbers from 1 to L3.
- Define one matrix of size L3 x nMoments, which we call meanMoments, and we initialize to contain all 0.
- Start an iterative process where iFShift goes from 1 to the size of frameshift and perform the following operations:
    - Declare frameshift as the value of frameshifts at iFShift;
    - Declare a matrix called moments of size nMoments x (L - frameshift), and we initialize it to contain all 0.
    - Define the index iMRow to iterate through moments rows.
    - Compute an array called dx (x distance) with (L - frameshift) elements that at each position contains the difference between the x value of points found at position iP and the x value of points found at position iP + frameshift.
    - Compute a corresponding array called dy (y distance),
    - Compute an array called d_sq (distance squared) that at each position contains [(dx^2 + dy^2) * micronPerPixel], where micronPerPixel is a parameter that defines the real-time pixel scaling of the imaging system under consideration. For artificial images (Rigano and Strambio-De-Castillia, 2018d), micronPerPixel is set to 1.
    - Start a nested iterative loop where iM_row goes from 1 to nMoments and perform the following:
        - Fill the iMRow row of the moments matrix with the d_sq array elevated at the (iMRow-1)/2 power.
    - Calculate the average value for each row of the moments array and save it to the column of the meanMoments matrix corresponding to the current value of iFShift.
- Compute delta_t as an array where position n is frameshifts at position n times parameter secPerFrame, which in case of artificial images we set to 1.
- In a loop where the index iMRow goes from 1 to nMoments, compute the following:
    - x as an array where the value at each position is computed as logarithm base 10 of the value of delta_t found at the same position.
    - y as an array where the value at each position is computed as logarithm base 10 of the value found in the meanMoments array at the position equal in value to iMRow.
    - c as the linear fit using x and y as coordinates
    - scalingCoefficients at the moment iMRow as the slope of the c linear fit
    - intercepts at the moment iMRow as the intercept of the c linear fit
- Compute mss as the array where position n is the scalingCoefficient at position n

- Compute D as the array where position n is 10 to the power of intercepts at position n divided by 4
- Compute x2 as a new array containing value from 0 to nMoments -1
- Compute y2 as a new array where position n is the value of mss at position n
- Compute c2 as a new array containing the linear fit obtained using x and y as coordinates
- Compute mssSlope as the slope of the c2 fit.
- Return D and mssSlope.

**calcDiffusion**

- Declare Px as an array containing the x-coordinates of the points of the input trajectory (e.g. from *BrownianTrajectoryGenerator*, *BrownianTrajectoryGenerator_2* or *TrajectoryGenerator*).
- Declare Py as the corresponding y-coordinates array.
- Define L as the length of trajectory
- Define minlag as the minimum size of the Δt time interval (i.e. window) used for subsequent computations and set 1 as the default value.
- Define maxlag as the maximum size of the Δt time interval and set L/2 as the default value (e.g. the values of maxlag employed in Supplemental Figures 5 were L/10 and L/250)
- Define no_overlap as a boolean where TRUE (i.e., 1) indicates the absence of overlap between subsequent computation windows, and the default value to 0.
- Define N as an array of whose length equals (maxlag-minlag) to be used to store the total number of datapoints used to calculate the squared displacement (SD) for each specific value of lag (i.e. frameshift)
- Define SD as an array of the same variable length as N to contain the total SD for each value of lag
- Define MSD as an array of the same variable length as N and SD that for each position will store either 0 or the corresponding value of SD divided by the corresponding value of N.
- Define di as an integer containing the current step
- In a for loop where lag goes from minlag to maxlag compute the following:
    - initialize N at position lag as zero
    - initialize SD at position lag as zero
    - define di as the step to be used in the computation, and initialize as lag if no_overlap = 1 otherwise as 1
    - in a for loop where i goes from lag to L with a step of di compute the following:
        - compute dx as (Px at point i) – (Px at point i - lag)
        - compute dy as (Py at point i )- (Py at point i - lag)
        - compute d2 as dx squared + dy squared
        - sum d2 to the current value of SD at position lag
        - increase the counter of the datapoint used and store in N at position lag

- if value of N at position lag is more than 0, compute MSD at position lag as SD at position lag divided by number of datapoint used else set MSD at position lag to zero

- Define ts as an array from minlag to maxlag
- Define linefit as a new linear fit using ts and MSD as coordinates
- Compute the intercept and slope of the linear fit
- Compute Dlin as the slope divided by 4
- Define log_linefit as a new linear fit using log_ts and log_MSD as coordinates
- Compute the intercept and slope of the linear fit
- Compute Dlog as the exp to the power of intercept divided by 4

#### Validation of *getMSS* against independent ODC estimation methods and using independently generated random walks

The functionality of the ODC and SMSS computation method implemented as part of the Diffusivity Measures plugin of OMEGA was compared with that of *getMSS* (Sbalzarini and Koumoutsakos, 2005; Racine, 2011). Because *getMSS* was never formally validated, the first step in this procedure consisted in testing the functionality of this algorithm on synthetic trajectories of known mobility (Supplemental Figures 5 and 6) and comparing the ODC and SMSS estimation results of *getMSS* with those obtained with independent motion type estimation methods. Because our *TrajectoryGenerator* Matlab routine (Helmuth et al., 2007; Rigano and Strambio-De-Castillia, 2018e; Rigano et al., 2018) depends on getMSS for the generation of trajectories of know mobility, we used the well-known random-walk simulation method to generate synthetic trajectories of known input D and SMSS (see above). While this method limits the scope of the analysis to trajectories that by definition will have a value of SMSS equal to 0.5, we reasoned this approach would be sufficient for our purposes.

Initially a limited test set of synthetic Brownian trajectories (input D = 0.01 and 0.1 µm2/s; input L = 1000, 10000, and 50000) were generated using the *BrownianTrajectoryGenerator* algorithms described above (Supplemental Figure 5, *BrownianTrajectoryGenerator*). Subsequently the ability of the log-log ODC estimation method of *getMSS* to correctly estimate D was assessed its results with the ground-truth as well as with results obtained using the linear and log-log ODC estimation methods implemented in *calcDiffusion* (Supplemental Figure 5, *log-log method* and *linear method*; see above). In all cases, two different calculation windows (i.e., L/25 and L/100, respectively) were used and relative distances from ground-truth (DGT) for D were computed as indicated:

$$Relative\ DGT_D = (D_{output} - D_{input})/D_{input}$$

As expected, the results of getMSS were indistinguishable from those obtained with the log-log method of *calcDiffusion* regardless of L, input D and calculation window (Supplemental Figure 5, *BrownianTrajectoryGenerator*). In addition, with small calculation windows (i.e., L/250) the log-log results were indistinguishable with results obtained with the linear D estimation method. Conversely, for larger computation windows (i.e., L/10) results produced using the *linear method* were significantly different both in terms of bias (i.e., mean DGT values) and variance of DGT from both log-log methods (Supplemental Figure 5). The most striking difference between *log-log* and *linear* methods was that while variance in the case of the linear method converged to a minimum value with length, in the case of either log-log methods variance appeared to increase for values of L > 1000. In order to ensure that this difference was not due to the synthetic random-walk generation algorithm employed in the first set of tests, a second set of test random walks was produced using an independent algorithm and showed the same results (Supplemental Figure 5, *BrownianTrajectoryGenerator_2*).

In order to extend the scope of *getMSS* validation, algorithm *BrownianTrajectoryGenerator* was utilized to produce sets of 1000 random walks to reflect 16 L x 11 D input values as indicated (Supplemental Table III). Values of ODC and SMSS were back-computed using *getMSS* with calculation windows of diminishing sizes (i.e., L/3, L/5 and L/10). Alongside $DGT_{ODC}$, relative SMSS DGT values were computed as follows:

$$Relative\ DGT_{SMSS} = (SMSS_{output} - SMSS_{input})/SMSS_{input}$$

and were plotted as a function of input ODC and L (Supplemental Figure 6). Consistent with results presented in Supplemental Figure 5, relative $DGT_D$ values showed a bi-modal distribution with trajectory length and no dependence on input D. Thus, DGT variances and, to a lesser extent, bias first converged towards zero and then climbed again for larger L values. Interestingly, and again consistent with Supplemental Figure 5, this effect diminished the shorter the calculation window.

In the case of SMSS, relative $DGT_{SMSS}$ values displayed a more canonical behavior with both bias values and variances of relative DGT converging to zero with increased length and with decreased window sizes, while again showing no dependence on input D.

#### Validation of OMEGA motion analysis routines

The performance of the OMEGA Diffusivity Measures plugin (Supplemental Figure 7) was validated by direct comparison with the previously validated *getMSS*. For this purpose, artificial trajectories were generated using the *TrajectoryGenerator* Matlab routine (Helmuth et al., 2007; Rigano and Strambio-De-Castillia, 2018e; Rigano et al., 2018) to reflect a total of 1936 test cases (Supplemental IIII). Values of SMSS were computed using both *getMSS* and OMEGA, Relative $DGT_{SMSS}$ were calculated, and the results compared (Supplemental Figure 7, top panel). As expected, results from the two algorithms were virtually indistinguishable, demonstrating the correctness of our Diffusivity Measures plugin implementation.

In the absence of a direct comparison with *getMSS*, relative $DGT_{ODC}$ results obtained with OMEGA were nonetheless plotted and their dependency on L, $ODC_{In}$ and $SMSS_{In}$ was evaluated (Supplemental Figure 7, bottom panel). While the overall error was found to be negligible and displayed the expected dependence on L, it was interesting to observe that it decreased with input SMSS and it instead increased with ODC.

### *Error propagation*

#### Calculation of the limits of detection for diffusivity

We employ the global error model described by Martin et al. (Martin et al., 2002; Rigano et al., 2018) to calculate Minimum Detectable ODC of order 2 ($ODC_{2\,MinDet}$) values as a function of image quality and detection offsets, as follows:

$$ODC_{2\,MinDet} = \frac{[bias_{PosV}(ImgMinSNR)]^2 + [\sigma_{PosV}(ImgMinSNR)]^2}{(2dt)}$$

Where: $Img_{MinSNR}$ is the minimum local SNR observed within the image under study; t is the time step; d is the space dimension (i.e., 2 for xy images; 3 for xyz images); PosV is the vector defining the position of a particle in Cartesian space; DGT denotes Distance from Ground Truth; $bias_{PosV}$ and $\sigma_{PosV}$ are measures of the particle detection error and imprecision, respectively (i.e., $bias_{PosV}$ equals the average of all observed values of $DGT_{PosV}$ = observed PosV - expected PosV; and $\sigma_{PosV}$ is their standard deviation; see above).

MOSAIC group. 2006. MOSAICsuite.

Object Refinery Inc. 2017. JFreeChart.

OME Consortium. 2017a. Bio-Formats.

OME Consortium. 2017b. OMERO.blitz.

Racine, R. 2011. Estimating the Hurst Exponent. I.F. Sbalzarini, editor. ETH Zurich, Zurich. 30 pp.

Rigano, A., and C. Strambio-De-Castillia. 2016a. OmegaProject/MIAPTE: repository. *github.com*. GPL–3.0–289. doi:https://doi.org/10.5281/zenodo.2535534.

Rigano, A., and C. Strambio-De-Castillia. 2016b. Minimum Information About Particle Tracking Experiments. 1st ed. Biosharing.org. 1 pp.

Rigano, A., and C. Strambio-De-Castillia. 2017a. OmegaProject/GenericTools: code repository. *github.com*. GPL–3.0. doi:https://doi.org/10.5281/zenodo.2538326.

Rigano, A., and C. Strambio-De-Castillia. 2017b. Proposal for minimum information guidelines to report and reproduce results of particle tracking and motion analysis. *bioRxiv*. 155036. doi:https://doi.org/10.1101/155036.

Rigano, A., and C. Strambio-De-Castillia. 2018a. OmegaProject/Omega: code repository. *github.com*. Java, C++, OMERO:GPL–3.0. doi:https://doi.org/10.5281/zenodo.2535523.

Rigano, A., and C. Strambio-De-Castillia. 2018b. OmegaProject/OmegaCommons: code repository. *github.com*. GPL–3.0. doi:https://doi.org/10.5281/zenodo.2538197.

Rigano, A., and C. Strambio-De-Castillia. 2018c. OmegaProject/OmegaOmeroCommons: code repository. *github.com*. GPL–3.0. doi:https://doi.org/10.5281/zenodo.2548938.

Rigano, A., and C. Strambio-De-Castillia. 2018d. OmegaProject/ArtificialImageGenerator: code repository. *github.com*. GPL–3.0. doi:https://doi.org/10.5281/zenodo.2555460.

Rigano, A., and C. Strambio-De-Castillia. 2018e. OmegaProject/TrajectoryGenerator: code repository. *github.com*. GPL–3.0. doi:https://doi.org/10.5281/zenodo.2555514.

Rigano, A., V. Galli, K. Gonciarz, I.F. Sbalzarini, and C. Strambio-De-Castillia. 2018. An algorithm-centric Monte Carlo method to empirically quantify motion type estimation uncertainty in single-particle tracking. *bioRxiv*. 379255. doi:10.1101/379255.

Ryan, T.A., P.J. Millard, and W.W. Webb. 1990. Imaging [Ca2+]i dynamics during signal transduction. *Cell Calcium*. 11:145–155.

Saxton, M.J. 2009. Single Particle Tracking. *In* Fundamental Concepts in Biophysics. Humana Press, Totowa,

NJ. 1–33.

Saxton, M.J., and K. Jacobson. 1997. Single-particle tracking: applications to membrane dynamics. *Annu Rev Biophys Biomol Struct*. 26:373–399. doi:10.1146/annurev.biophys.26.1.373.

Sbalzarini, I.F., and P. Koumoutsakos. 2005. Feature point tracking and trajectory analysis for video imaging in cell biology. *J Struct Biol*. 151:182–195. doi:10.1016/j.jsb.2005.06.002.

SciJava consortium. 2017. SciJava.

SQL project. 2018. MySQL Connector/J.

SWT consortium. 2018. SWT: The Standard Widget Toolkit.

Xiao, X., V.F. Geyer, H. Bowne-Anderson, J. Howard, and I.F. Sbalzarini. 2016. Automatic optimal filament segmentation with sub-pixel accuracy using generalized linear models and B-spline level-sets. *Medical Image Analysis*. 32:157–172. doi:10.1016/j.media.2016.03.007.
