## Supplementary material for "OMEGA: a software tool for the management, analysis, and dissemination of intracellular trafficking data that incorporates motion type classification and quality control": 02_Supplemental Information 2

# getMSS pseudocode
# getMSS is a Matlab function developed to estimate the Observed Diffusion Constant (ODC) # and the Slope of Moment Scaling Spectrum of existing trajectories 
# (Sbalzarini & Koumoutsakos, 2005; Racine, 2011) 

Points = paired (x, y) coordinates of points from input trajectories; 
nMoments = number of moments of displacement;
L = length of trajectory;
L3 = L / parameter that define the size of windows;
frameshifts = array from that contains number from 1 to L3;
meanMoments = matrix of size nMoments x L3 filled with zeros;
for iFShift from 1 to size of frameshifts
➢	frameshift = frameshifts at iFShift;
➢	moments = matrix of size nMoments x (L - frameshift) filled with zeros;
➢	dx = x points(from 1 to L-frameshift) -  x points(from 1+frameshift to L);
➢	dy = y points(from 1 to L-frameshift) -  y points(from 1+frameshift to L);
➢	d_sq (pos n) = (dx (pos n) * dx (pos x) + dy (pos n) * dy pos(x))*micronPerPixel*micronPerPixel;
		•	for iMRow from 1 to nMoments
		•	moments(iMRow row) (pos n) = d_sq (pos n) ^((iMRow-1)/2);
		•	end for
➢	meanMoments( iFShift column ) (pos n) = average of moments of displacement n row 
end for
delta_t (pos n) = frameshifts (pos n) * secPerFrame;
for iMRow from 1 to nMoments
➢	x (pos n) = log10(delta_t (pos n));
➢	y (pos n) = log10(meanMoments(iMRow row pos n));
➢	c = polyfit(x,y,1);
➢	scalingCoefficients(iMRow) = c(1);
➢	intercepts(iMRow) = c(2);
end for
mss = scalingCoefficients;
D = (10.^(intercepts))/4;
x2 = 0:(nMoments-1);
y2 = mss;
c2 = polyfit(x2, y2, 1);
mssSlope = C2(1);
return D and mssSlope

#*****************************************************************************
# this file is released under a Creative Commons Attribution Share Alike 4.0 International # License (https://creativecommons.org/licenses/by-sa/4.0/)
