## Supplementary material for "OMEGA: a software tool for the management, analysis, and dissemination of intracellular trafficking data that incorporates motion type classification and quality control": 03_Supplemental Information 3

#calcDiffusion pseudocode
# calcDiffusion is a Perl script developed to estimate the Observed Diffusion Constant #(ODC) of existing trajectories 
# Lawrence Lifshitz, 2015 unpublished

for lag from minlag to maxlag included
➢	N[lag] = 0;						# number of data points for this lag
➢	SD[lag] = 0;					# total squared displacement for this lag
➢	MSD[lag] = 0;					# mean square displacement for this lag
➢	if no_overlap then di = lag else di = 1;
➢	for i from lag to L with step of di
		•	dx = Px [i] - Px [i-lag];
		•	dy = Py [i] - Py [i-lag];
		•	d2 = dx*dx + dy*dy;
		•	SD[lag] += d2;
		•	N[lag]++;				# count of how many data points
➢	end for
➢	if N[lag] then MSD[lag] = SD[lag]/N[lag]  else MSD[lag] = 0;
end for
ts = (minlag .. maxlag);
lineFit = Statistics::LineFit->new();
lineFit->setData(ts,MSD);
(intercept, slope) = lineFit->coefficients();
Dlin = slope/4;
log_lineFit = Statistics::LineFit->new();
log_ts = array_log(ts);  
log_nMSD = array_log(MSD);
log_lineFit->setData(log_ts, log_nMSD);
(log_intercept, log_slope) = log_lineFit->coefficients(); 
Dlog = exp(log_intercept)/4;

#*****************************************************************************
# this file is released under a Creative Commons Attribution Share Alike 4.0 International # License (https://creativecommons.org/licenses/by-sa/4.0/)
