## Supplementary material for "OMEGA: a software tool for the management, analysis, and dissemination of intracellular trafficking data that incorporates motion type classification and quality control": 04_Supplemental Figures

### Table of content

**Supplemental Figure 1:** **Typical OMEGA dataflows**

**Supplemental Figure 2**: **OMEGA for developers: logical structure and software architecture**

**Supplemental Figure 3**: **Uniform artificial trajectories examples**

**Supplemental Figure 4**: **Validation of the SNR Estimation plugin**

**Supplemental Figure 5**: **Comparison of getMSS with alternative ODC estimation methods**

**Supplemental Figure 6**: **Validation of getMSS over extended Brownian trajectories test sets**

**Supplemental Figure 7**: **Validation of OMEGA motion type estimation**

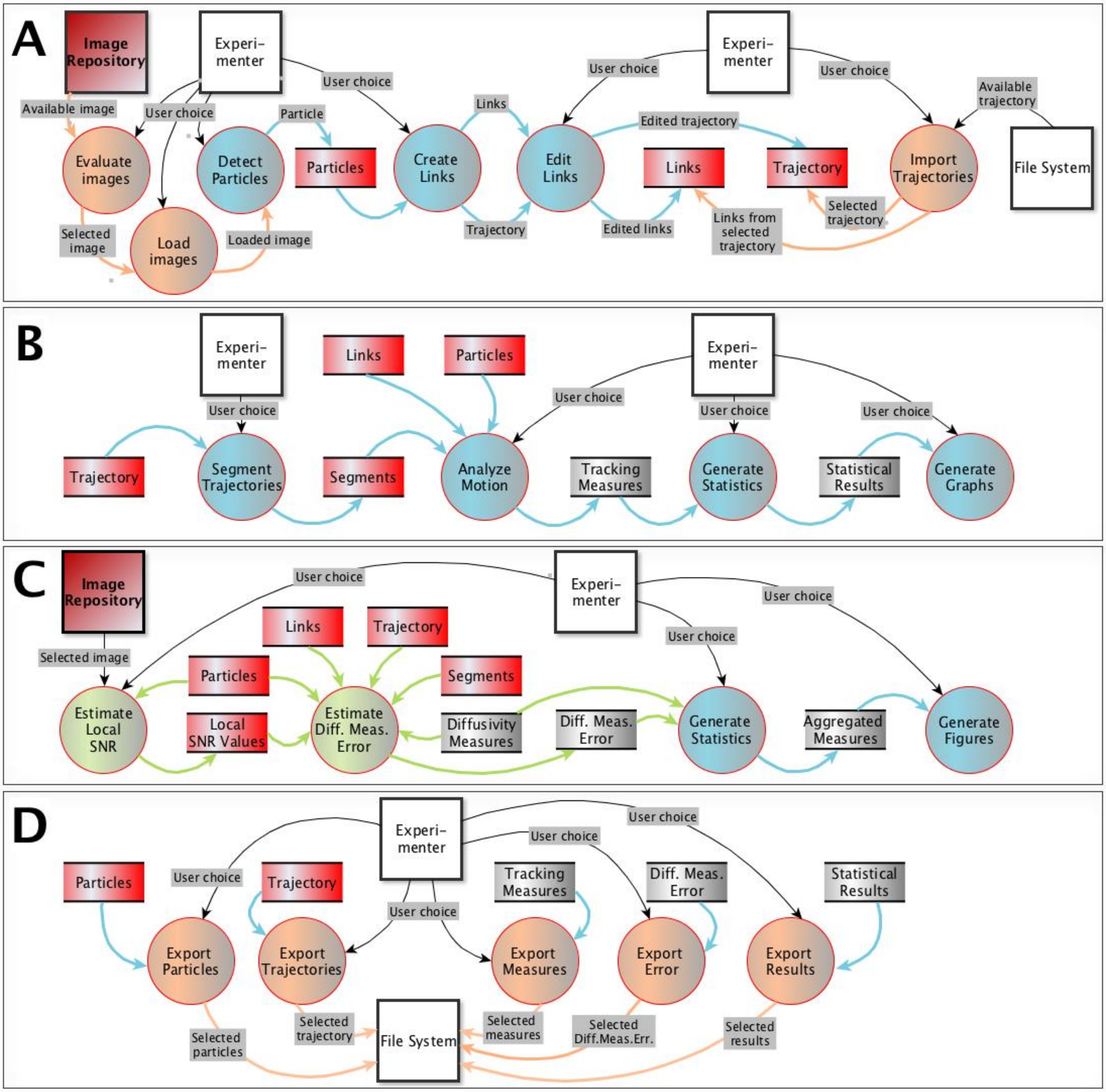


### *Supplemental Figure 1:*

**Typical OMEGA dataflows.** Four common paths data take in OMEGA three are depicted here using the data flow diagram (DFD) formalism. Circles represent processes that transform data, arrows represent data in motion, labels on arrows represent specific data packages being moved, double lined-rectangles represent data at rest (i.e. data stores) and squares represent entities (i.e. Experimenter(s) and external data repositories) that interact with the system from the outside. A) Images selection, tracking and import of pre-computed particles and trajectories from the file system. B) Trajectory editing, segmentation, and motion analysis. C) Estimation of error associated with diffusivity measures. Labels are often omitted for simplicity sake. D) Export of identified particles, trajectories, tracking measures, diffusivity measures errors and statistical results to the file system.

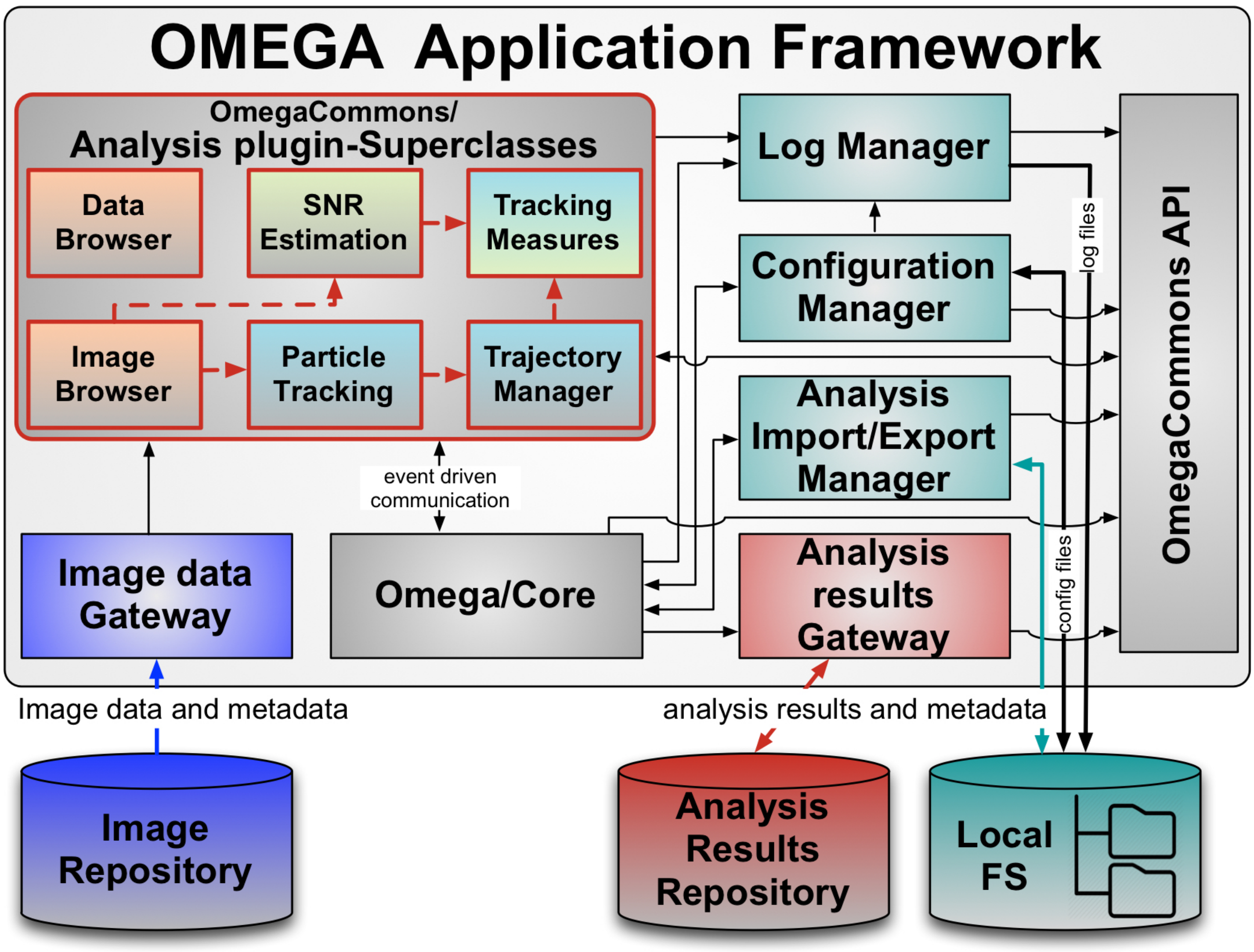


### *Supplemental Figure 2:*

**OMEGA for developers: logical structure and software architecture.** The OMEGA application is composed of three core structural elements (*grey boxes*) that work in concert with five input/output components (*colored boxes*) to communicate with external data stores consisting of a third party Image Repository (*blue cylinder*), the dedicated OMEGA Analysis Results Repository (*red cylinder*) and the File system (*green cylinder*). The three structural components are the: Omega/*Core, OmegaCommons/Analysis plugin-Superclasses,* and *OmegaCommons API. Omega/Core* contains all essential sub-components needed by OMEGA to launch and run, including the application launcher, the initialization and disposition methods, and the event-driven logic driving all communication between the core and *Analysis plugin-Superclasses.* The latter consists of six modular super-classes that contain all the main functional logic of OMEGA and are extended by interchangeable plugins (Figure 3) to carry out the depicted (dashed red arrows) data-flow. Color coding for *Analysis plugin-Superclasses* is as Table I and Figure 3. *OmegaCommons API* was designed to facilitate the extraction of OMEGA libraries and aide custom plugin development. In addition to general utility methods, it includes the Java building blocks of individual plugins (including all foundational libraries necessary for trajectory analysis), of interfaces necessary for the communication between plugins and the Omega/*Core*, and of the event-driven communication system. *OmegaCommons API* also contains all foundational libraries necessary for trajectory analysis in OMEGA. The five communication components comprise the *Log*, *Configuration* and *Analysis Import/Export Managers* that interact with the FS, as well as two *Gateways* that are required to execute all necessary input-output tasks between the application and the data repositories.

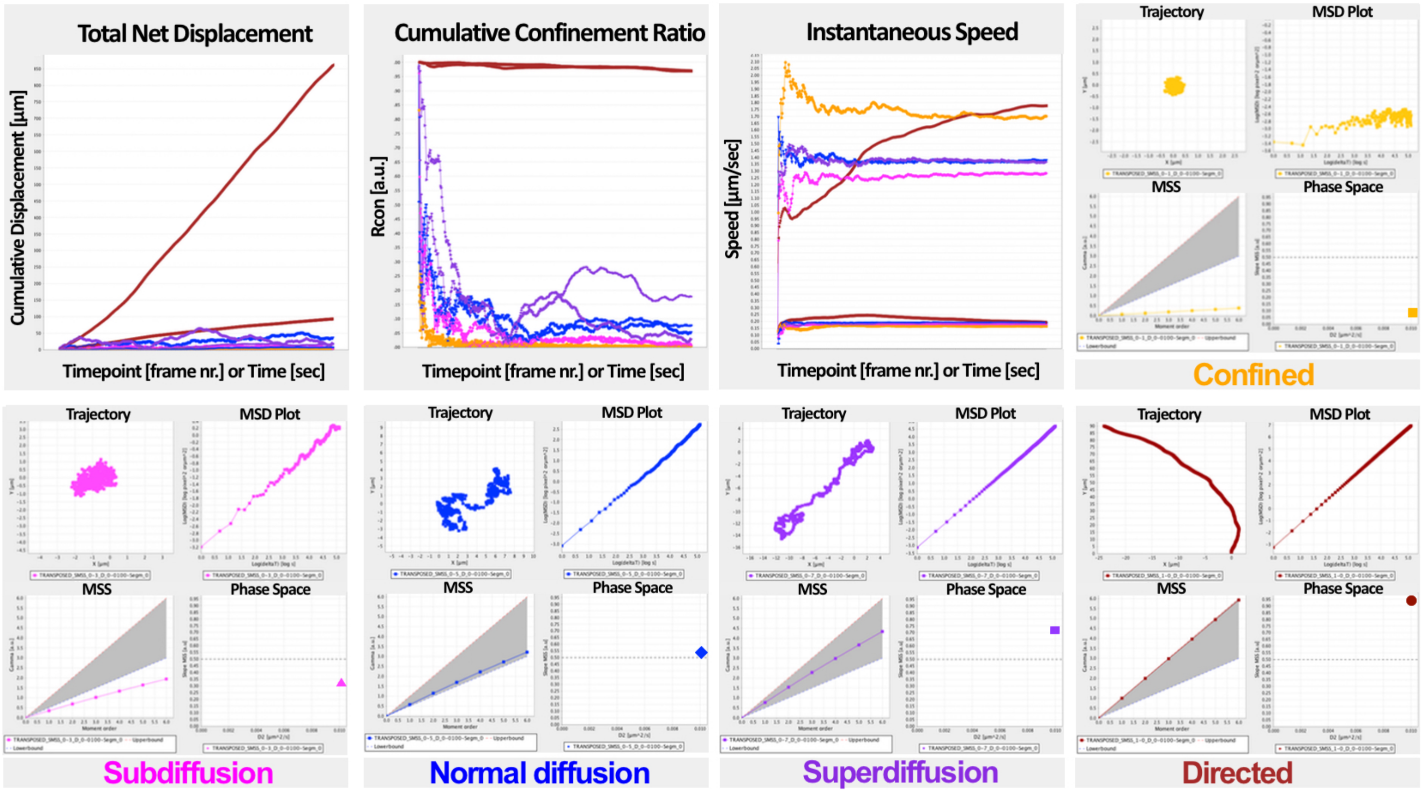


### *Supplemental Figure 3:*

**Uniform artificial trajectories examples.** Ten self-similar artificial trajectories of known mobility were generated using our *TrajectoryGenerator* MatLab algorithm as described (Figure 5; Supplemental Information 1). After importing into OMEGA, they were assigned the corresponding motion type label by using the Trajectory Segmentation plugin and then subjected to motion analysis using the Mobility, Velocity and Diffusivity Tracking Measures plugins. Top row (left to right): 1) plot displaying the total net displacement of each of the ten artificial trajectories as a function of time; 2) plot displaying the cumulative confinement ratio of each of the ten artificial trajectories as a function of time; 3) plot displaying the instantaneous speed of each of the ten artificial trajectories as a function of time; and 4) motion-type classification plot set for artificial trajectory of confined motion type (ODC = 0.01; SMSS = 0.1). Bottom row (left to right): 1) motion-type classification plot set for artificial trajectory of sub-diffusive motion type (ODC = 0.01; SMSS = 0.3); 2) motion-type classification plot set for artificial trajectory of normal diffusive motion type (ODC = 0.01; SMSS = 0.5); 3) motion-type classification plot set for artificial trajectory of super-diffusive motion type (ODC = 0.01; SMSS = 0.7); and 4) motion-type classification plot set for artificial trajectory of directed motion type (ODC = 0.01; SMSS = 1.0).

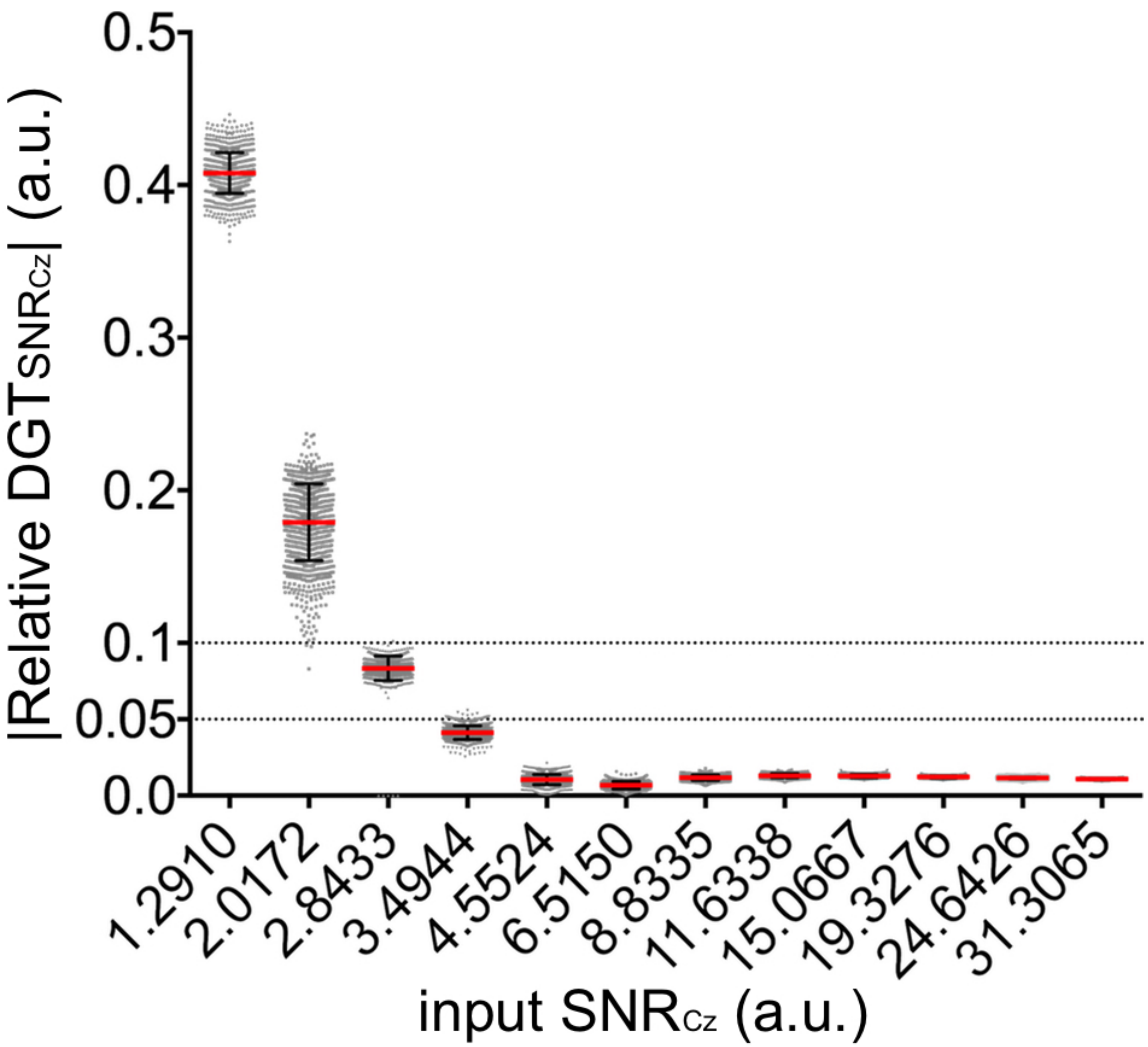


***Supplemental Figure 4:***

**Validation of the SNR Estimation plugin.** The ability of the OMEGA implementation of the *MOSAICsuite*'s local SNR Estimation algorithm (Sbalzarini & Koumoutsakos, 2005; Xiao *et al*, 2016; Gong & Sbalzarini, 2016) to correctly estimate SNR according to Cheezum (Cheezum *et al*, 2001) was assessed on artificial images presenting moving point sources with varying input local SNR characteristics. Distributions of absolute values of relative distances from ground truth (DGT) between input and output SNR values ($|Relative\ DGT_{SNR_{Cz}}|$) were computed and are presented here as swarm plots as a function of input SNR. Red lines indicates the mean value of each distribution and black lines represent the +/- 1 x standard deviation interval around the mean (i.e. 68% confidence interval).

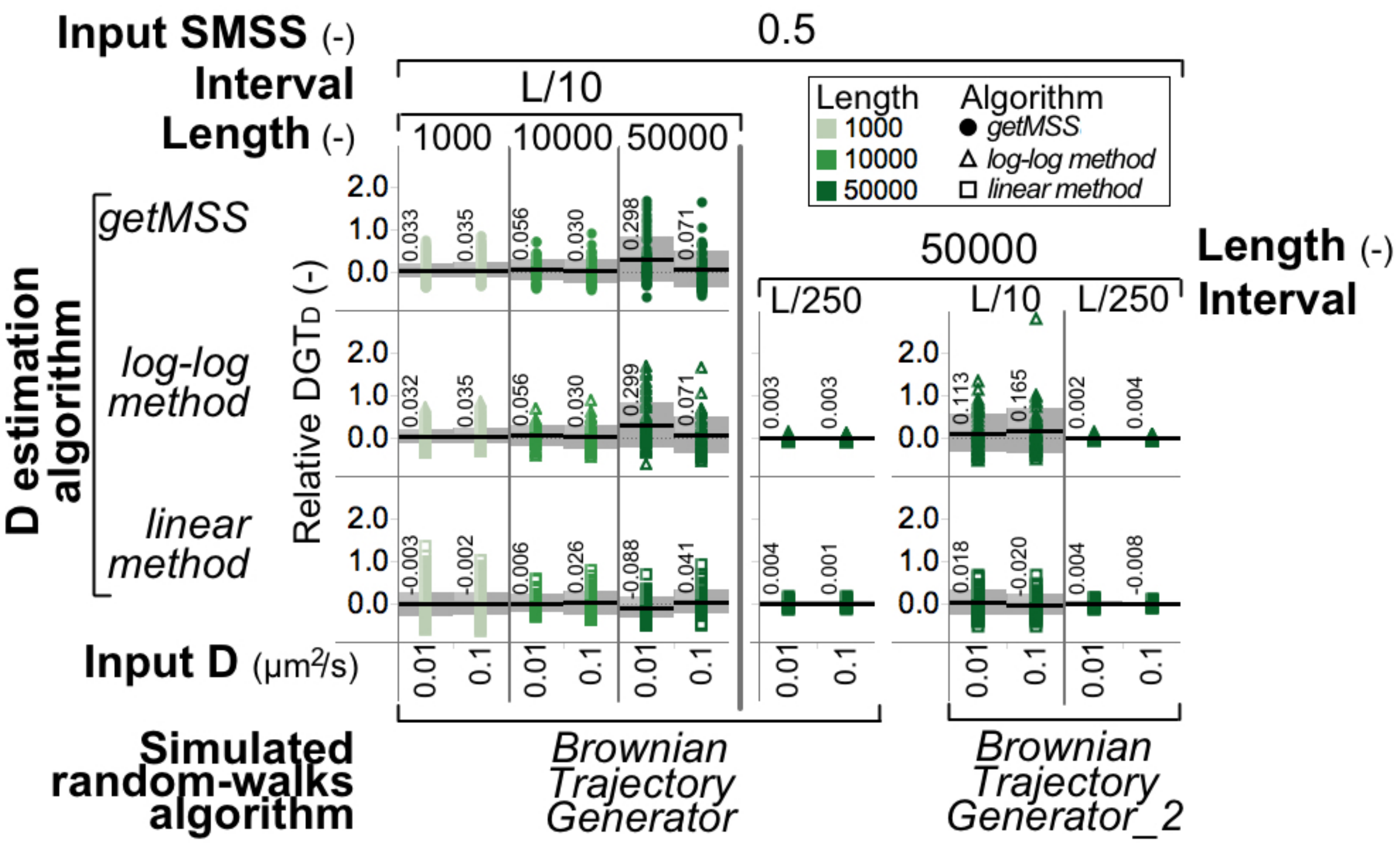


***Supplemental Figure 5:***

**Comparison of getMSS with alternative ODC estimation methods.** The ability of the Matlab *getMSS* motion type estimation routine (Supplemental Information 1) to correctly compute ODC values was assessed by comparing its performance with that of the two ODC estimation methods found in the Perl *calcDiffusion* algorithm (*log-log method* and *linear method*), on artificial Brownian trajectories produced using two independent methods for random-walk generation (*BrownianTrajectoryGenerator* and *BrownianTrajectoryGenerator_2*). Trajectories were generated using 3 input L times 2 input D values (i.e., for a total of 6 test cases; Supplemental Table III). D estimation was performed using Δt intervals (*Interval*) equal to L/3 and L/5 and L/10, as indicated. Results are presented as scatter distributions of individual relative Distance from Ground Truth for D ($DGT_D$) values. Black lines and associated numbers represent average $DGT_D$ values and grey areas indicate the +/- 1 x standard deviation interval around the mean. D values are calculated as µm^2/s assuming 1 pixel/µm and 1 frame/s. In all cases, n = 1000 artificial trajectories per test set.

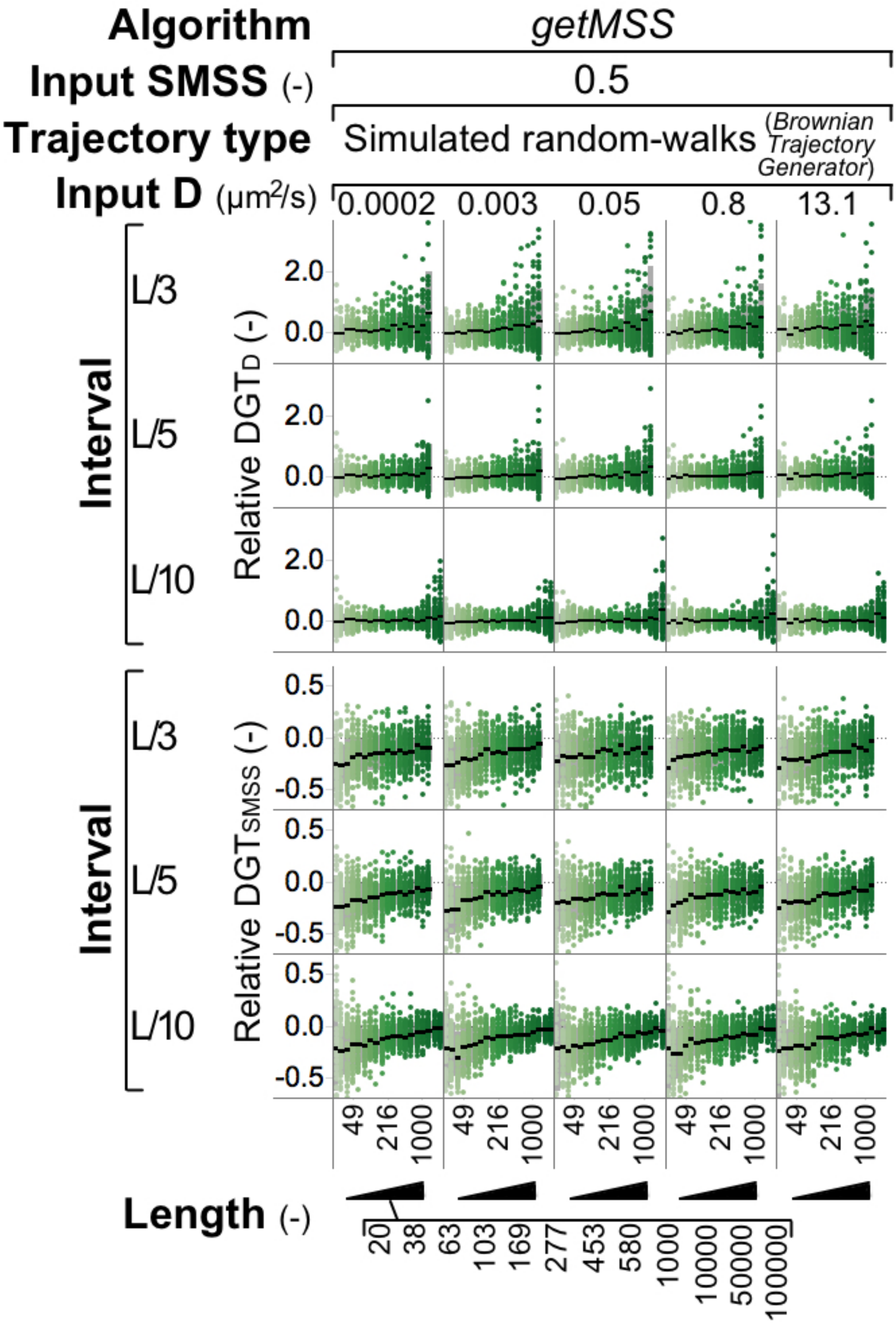

Algorithm
getMSS
Input SMSS (-)
0.5
Trajectory type
Simulated random-walks (Brownian Trajectory Generator)
Input D (µm²/s)
0.0002
0.003
0.05
0.8
13.1
Interval
L/3
L/5
L/10
Relative DGTD (-)
2.0
0.0
Relative DGTSMSS (-)
0.5
0.0
-0.5
49
216
1000
Length (-)
20
38
63
103
169
277
453
580
1000
10000
50000
100000

### *Supplemental Figure 6:*

**Validation of getMSS over extended Brownian trajectories test sets.** The validation of *getMSS* (Supplemental Information 1) was extended by evaluating its ability to correctly compute D and SMSS on an extended test set of random-walks artificially generated using *BrownianTrajectoryGenerator* (Supplemental Information 1). Trajectories were generated using 19 input L times 11 input D values (i.e., for a total of 209 test cases; Supplemental Table III) of which only a subset are shown as indicated. D and SMSS estimation was performed using Δt intervals (*Interval*) equal to L/3 and L/5 and L/10, as indicated. Results are presented as scatter distributions of the observed values of relative Distance from Ground Truth for D ($DGT_D$) and for SMSS ($DGT_{SMSS}$). Black lines represent average DGT values and grey areas indicate the +/- 1 x standard deviation interval around the mean. D values are calculated as µm^2/s assuming 1 pixel/µm and 1 frame/s. In all cases, n = 1000 artificial trajectories per test set.

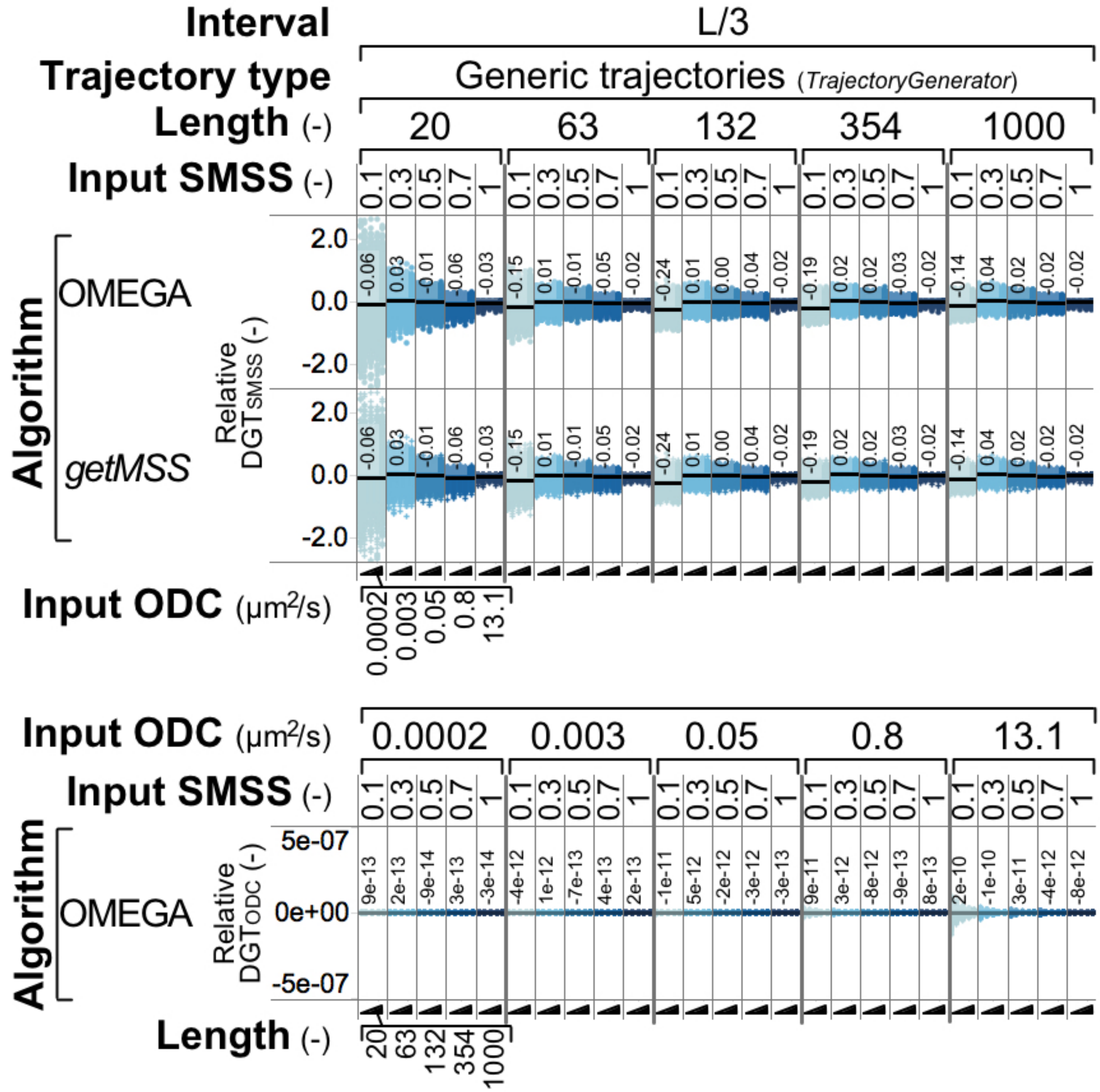


***Supplemental Figure 7:***

**Validation of OMEGA motion type estimation.**

The ability of the OMEGA Diffusivity Trajectory Measures plugin to correctly compute SMSS (Top) and ODC (Bottom) was assessed by comparing its performance with that of *getMSS*. *TrajectoryGenerator* (Supplemental Information 1) was used to generate 1936 test sets of artificial trajectories (Supplemental Table III) of which only a subset are shown. SMSS and ODC estimation was performed using Δt intervals (*Interval*) equal to L/3 and L/5 and L/10, as indicated. In all cases input SMSS and ODC values were compared with the corresponding output values and plotted as a function of L, SMSS and ODCD. Results are presented as scatter distributions of the observed values of relative Distance from Ground Truth for D ($DGT_{ODC}$) and for SMSS ($DGT_{SMSS}$). Black lines and associated numbers represent average DGT values and grey areas indicate the +/- 1 x standard deviation interval around the mean. ODC values are calculated as µm^2/s assuming 1 pixel/µm and 1 frame/s. In all cases, n = 1000 artificial trajectories per test set.
